# A scalable genomic framework for programmable strain tagging in a diverse bacterial genus

**DOI:** 10.64898/2026.02.24.707161

**Authors:** Ezgi Mehmetoğlu Boz, Hugo R. Barajas, Tsung-Ta Yu, Mounika Thigulla, Naila Nazir, Kyle A. Gervers, Sarfaraz Hussain, Lorena Carot Hernández, Derek S. Lundberg

## Abstract

Transposons are a convenient vehicle for inserting DNA into a bacterial genome, but widely-used transposons such as Tn7, Tn5, and Mariner do not hit custom targets. Recently, CRISPR-associated transposons (CASTs) have been developed as tools to direct the insertion of a transposon to a chosen site with a short guide sequence. We adapted this system for use in the widespread plant-associated genus *Sphingomonas*, uniquely tagging genetically diverse strains in several different sites, and we demonstrate the utility of the tags for quantitative strain tracking in complex bacterial populations. Although we initially targeted the conserved site into which Tn7 integrates as a benign transposition location, a genomics search revealed insertions at this site would frequently disrupt genes not only in *Sphingomonas* but also in widely studied *Pseudomonas*. Therefore we identified improved genus-wide conserved sites between convergently terminating genes as alternatives. We experimentally validated an improved neutral site in *Sphingomonas*, and then targeted this site in a heterogeneous uncharacterized *Sphingomonas* population. Using a tagIM seq, a novel rapid transposon mapping method introduced here, we screened resulting transformant colonies for off-target transposon insertions. This enabled prioritization of correctly tagged novel strains, accelerating the creation of fully tagged synthetic communities for the high-throughput study of fine-scale bacterial natural variation.

## Introduction

To study an organism in its environment, one must avoid mistaking similar “look-alikes” for the organism of interest. Yet, distinguishing one organism from another can be challenging, especially for bacteria that often grow in complex, heterogenous communities. While bacteria can sometimes be identified using DNA sequencing, innate genetic differences are not helpful when individuals are genetically very similar or even identical to others in the population. Consequently, experiments relying on innate genetic differences must be carefully controlled to avoid external contamination, and all bacteria used in such experiments should be distinguishable from each other. This limits community size and environmental complexity [1].

As an alternative to detecting subtle innate features, a common approach is to modify bacteria with a unique element such as a fluorescent protein, an antibiotic resistance marker, and/or a unique DNA sequence that serves as a barcode. This approach of “tagging” bacteria, particularly through the use of antibiotic resistance and fluorescence, has been integral to the study of host-microbe interactions [2–4], and has allowed detailed characterization of individual genotypes in complex environments. While unique antibiotic resistances and fluorescence proteins are limited in variety [5], DNA barcodes can distinguish myriad individuals when detected and quantified by modern high-throughput sequencing methods, enabling the rigorous study of increasingly multipartite interorganismal interactions.

A major challenge for all tagging strategies is physically introducing the tags. Although genetic modification of some organisms like *E.coli* has become very efficient [6], the tools available to insert DNA into non-model organisms are more limited. Transposons are an especially attractive tool because they retain function in diverse bacteria without needing to be customized for each individual. Further, they can carry large payloads involving thousands of base pairs, and can integrate into the chromosome, making them less transient than broad host range plasmids which can be lost [7–9]. In particular, the Tn7 transposon is popular for high-throughput tagging strategies [7,8,10] because of its broad host range in Gram-negative bacteria and predictable recognition of the chromosomal attTn7 locus leading to insertion just downstream of the Glutamine--fructose-6-phosphate aminotransferase (*glmS*) gene. In most cases, Tn7 integrations are not associated with fitness penalties [8,11].

Like Tn7, CRISPR RNA-guided integrases, including the type I-F *Vibrio cholerae* CRISPR-associated transposons (*Vch*CAST) Tn6677-like elements [9,12], are functional in diverse bacteria. Unlike Tn7, they have the major advantage that the integration site can be programmed using a guide RNA comprising a 5’ CN PAM site immediately followed by a 32 bp sequence, with the transposon usually integrating ∼50 bp downstream of the guide sequence. With CASTs, it therefore possible to not only insert a transposon into an intergenic neutral site, but also to target and disrupt genes. In this work, we adapted a single vector *Vch*CAST system [9] for broad use in diverse strains of the widespread plant-associated bacteria *Sphingomonas*, ubiquitous Gram-negative Alphaproteobacteria that often grow in highly heterogeneous communities, and which are of interest for beneficial features including plant growth promotion, xenobiotic degradation, and industrial production of extracellular polymers [13–16].

We created a minimalist tagging construct including both conserved 16S rRNA gene (hereafter rDNA) priming sites and unique priming sites to amplify the DNA barcodes, and used genomic methods to identify conserved locations for versatile guide sequences that would enable targeting multiple diverse isolates. We then tagged five *Sphingomonas* strains in at least two genomic locations, revealing by extensive characterization of these insertions the flexibilities and limitations of the system. In controlled experiments with synthetic and wild communities, we validated the tags for quantitative strain tracking along with the background microbiota in a complex host-associated environment, identifying tag-specific amplification biases and an empirical method for determining correction factors. Importantly, our genus-wide analyses suggest that site into which Tn7 integrates is not only unlikely to be a safe integration site in many *Sphingomonas* strains, but also in other widely-studied bacteria including *Pseudomonas*. As an alternative to the Tn7 insertion location, we identify improved highly conserved sites for benign transposon integration in both *Sphingomonas* and *Pseudomonas*, and successfully target one such site in *Sphingomonas*. Further taking advantage of its conservation, we designed a tandem array of three related guide RNAs to broadly target this site and used the array to directly transform a heterogenous and uncharacterized *Sphingomonas* population.

Using tagIMseq, a novel rapid method for transposon mapping we introduce here, we identified colonies of previously uncharacterized strains that were correctly and uniquely barcoded, representing a shortcut that can be exploited to introduce unique DNA barcodes into large diverse populations for large-scale experiments investigating natural bacterial genetic variation.

## Results

### Design of a minimalist mutation and tagging construct

We designed a DNA barcoding construct (**Fig 1A**, **Supplementary Fig 1**) for the smallest footprint and lowest metabolic cost. As an antibiotic resistance marker necessary for efficient selection of transformed cells, we chose tetracycline resistance regulated by the native *tet*R repressor [17]. This system is broadly effective in *Sphingomonas* [18] including in strains from our collection, and importantly resistance is not actively transcribed in the absence of tetracycline and is therefore off in most experimental settings [19]. We flanked the tetracycline resistance gene with flippase (*frt*) sites that would allow it to be fully excised if desired, and did not include fluorescence proteins or any other constitutively expressed genes because these have limited uniqueness and may add metabolic costs. Downstream of the tetracycline resistance cassette we included two *rrnB* rho-independent transcriptional terminators to prevent transcription beyond the insertion. Following the terminator region is a unique 16 bp random sequence serving as a DNA barcode. The terminator and barcode are flanked by PCR priming sites, including not only unique priming sites but also the common 16S rRNA gene (rDNA) priming sites 515F and 799R/806R that amplify variable region 4 (V4) [20]. The full amplifiable region is ∼300 bp, nearly identical in length to actual bacterial V4 16S rDNA amplicons and long enough to be purified efficiently with Solid Phase Reversible Immobilization (SPRI) beads. We modified the pSPIN pSL1142 vector from [9] to include this barcoding construct on the transposon (**Methods**), and further modified the vector backbone to include a dominant negative synthetic *rpsL* gene that could confer streptomycin sensitivity, and thus be used for negative selection in *Sphingomonas* which are broadly naturally resistant to streptomycin [18]. The final vector library, loaded with random barcodes, we named pSPIN-LL.

**Figure 1.**
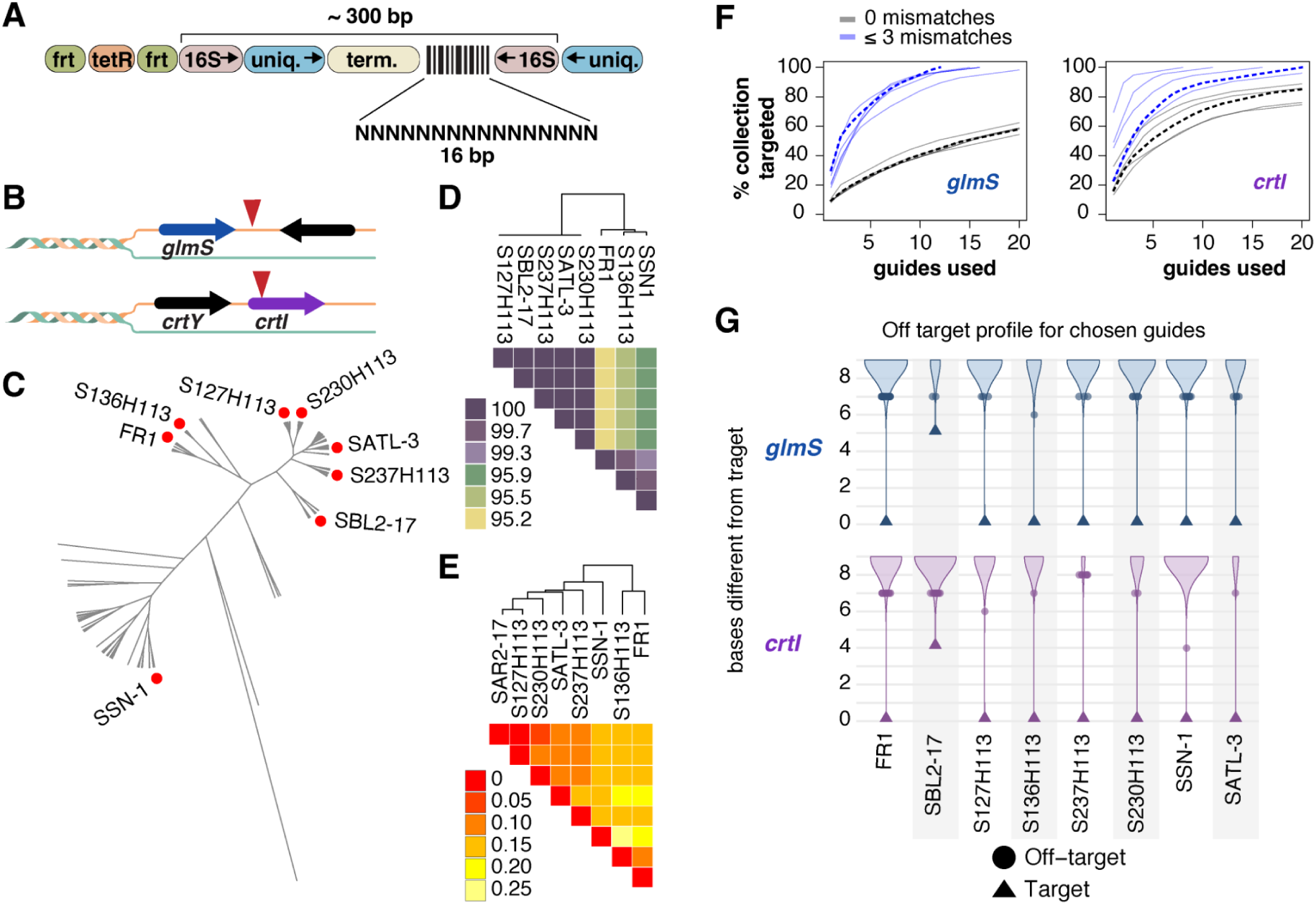
Design and planning of mutation constructs. **A)** Design of transposon payload for pSPIN-LL. frt, flippase sites; tetR, tetracycline resistance; uniq., unique priming sites; 16S, 16S rDNA priming sites; N, random nucleotides; term., terminators. Arrows represent the direction of primer extension during PCR. **B)** Diagram indicating targeted region for benign genome tagging (top, *glmS*) and for gene disruption (bottom, *crtI*). **C)** Unrooted phylogenetic tree of 138 nonredundant *Sphingomonas* genomes, with positions of transformed strains indicated by a red dot. **D**) Comparison of nucleotide identity in the V4 16S rDNA sequences of the transformed strains. **E)** Comparison of MASH distance between whole genome sequences of the transformed strains. **F)** Percentage of strains from our culture collection (y-axis) that can in principle be cumulatively targeted by a family of up to 20 guide sequences (x-axis). Each grey or blue line represents a family of 20 sequence-related guides derived from a single conserved PAM site, with grey lines representing the case of perfect alignment to the culture collection, and blue lines allowing for alignments with up to 3 mismatches. The dotted line represents the guide family used in experiments. **G)** Sequence similarity of off-target regions in each genome to the guide sequence designed for that genome as determined via BOWTIE2 alignment. Circles are plotted to represent discrete off-target events only for the most similar off-targets.

### Site-specific integration in diverse *Sphingomonas* strains

We chose 8 *Sphingomonas* strains from our sequenced culture collection that represented diverse lineages (**Fig 1C-E**) to gauge broad applicability in the genus, and prepared to transform each in both a neutral location and within a coding sequence (**Fig 1B**). For the neutral location, we initially reasoned that the Tn7 integration site about 25 bp downstream of the *glmS* stop codon would be ideal, as it is widely used and negative fitness effects are rarely reported. Since *Vch*CASTs insert around 50 bp downstream of the guide sequence, we only considered guide sequences within the final 75 bp of *glmS*, which should all lead to insertions after the stop codon. We looked for a site with minimal variation between all strains in our collection so that each guide would have utility in multiple strains, and identified six sites in the alignment that had 5’-CN-3’ PAM motifs in at least 97% of strains. For each candidate site, we asked which fraction of our culture collection could be targeted with a limited number of guides, and discovered that a family of 10 guides would cover about 40% of the collection assuming perfect matches, and around 90% if allowing for 3 mismatches (**Fig 1F**). Initially we assumed perfect matches may be important, and arbitrarily chose one of the guide families with which to proceed (**Fig 1F**, dotted line). We then checked that each guide in that family was unique within its target genome (**Fig 1G**). Due to a serendipitous error discovered later, the guide we designed for SphUPP:SBL2-17 differed from its actual genomic target by 5 bp (**Fig 1G, Supplementary Fig 2**).

As a coding sequence to disrupt, we targeted the phytoene desaturase (*crtI*) gene (**Fig 1B**), which is a critical upstream step in the production of the carotenoids that give *Sphingomonas* their vivid yellow and orange colors [21]. Mutations in *crtI* provide a clear white colony phenotype that allows successful mutagenesis to be assessed without molecular methods [18]. We targeted conserved PAM sites in the first 100 bp of the coding sequence and found five sites conserved in 97% of strains. A family of ten guides could target over 60% of the collection assuming perfect matches, and up to 100% allowing for 3 mismatches (**Fig 1F**). As with *glmS*, we chose a guide family and confirmed its uniqueness within each target genome (**Fig 1G**). For *crtI* also, the guide for SphUPP:SBL2-17 differed from its actual genomic target (**Fig 1G**, **Supplementary Fig 2**).

We synthesized all chosen guides (**Supplementary Table 1 and 2**), ligated each into pSPIN-LL, and then conjugated pSPIN-LL into the appropriate *Sphingomonas* (**Methods**). We then plated on tetracycline plus streptomycin to select for transformants that had lost the pSPIN-LL plasmid. For both *glmS* and *crtI* mutants, we performed two colony PCRs: one to verify that the barcoded tagging construct was present, and a second to verify that the pSPIN-LL plasmid could not be detected. We recovered at least one transformant in all 8 strains, demonstrating that integration is broadly possible in the genus. For subsequent analysis, we decided to focus on those five strains for which we recovered integrations resulting from both *crtI* and *glmS* guides. We attempted to map the insertion site(s) using transposon site insertion sequencing, but this was initially unsuccessful due to factors we subsequently optimized (see below), so for expediency we selected a single candidate colony from each transformation and determined the transposon’s exact location and barcode sequence using pooled whole genome sequencing with long reads (**Supplementary Table 3**, **Fig 2A-B**).

**Figure 2.**
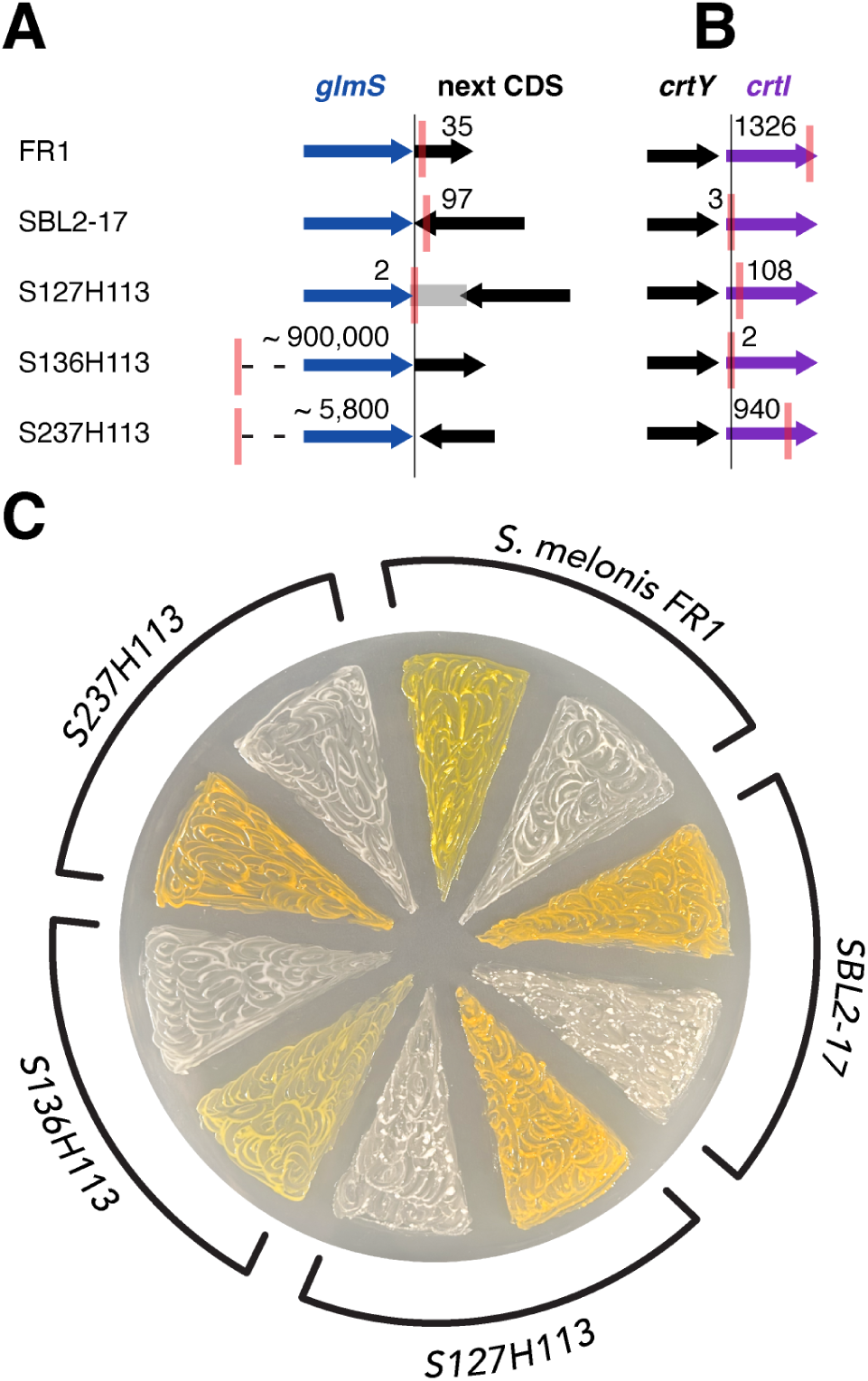
Tagging construct insertion locations. **A)** Diagram of mapped transposon insertions resulting from guides targeting the region downstream of *glmS*. The thin vertical line intersecting all rows represents the expected insertion location 50 bp downstream of the guide. while the red vertical line within each row represents the observed location. The calculated distance in base pairs between the observed and expected location is indicated by an integer. The grey box for SphATUE:S127H113 (*glmS*:Tn) represents deleted bases. CDS, coding sequence. **B)** Same as (A), for insertions in *crtI*. **C)** The *glmS*-targeting and *crtI*-targeting (pale color) transformants from each strain.

For all ten insertions, the transposon was present as a single insertion in the forward orientation (**Supplementary Fig 1**). For the five *crtI* mutants, the insertion occurred in the coding sequence and all transformants had lost their pigment (**Fig 2C**). For three of the *glmS*-targeting strains, the insertion occurred fewer than 200 bp downstream of the guide, while in the other two the insertion occurred thousands of bases away (**Fig 2A**). In SphATUE:S127H113, transposition created a deletion of 700 bp such that the final bases of the next ORF were eliminated (**Fig 2A**). Remarkably, both the *glmS* and *crtI*-targeting insertions in SphUPP: SBL2-17 were within 100 bp of their target despite the 5 or 4 respective mismatches in the guide, demonstrating that 85% nucleotide identity is sufficient if the target is unique!

### Fitness of tagged strains relative to untagged counterparts

While mapping the insertions, we increasingly questioned whether the Tn7 insertion site was a safe choice in *Sphingomonas*. Ideally, the transposon should insert between two convergently terminating genes in a region free of annotated features to avoid disrupting operons or regulatory elements, but in only three of our five transformed strains was *glmS* convergently terminating with the next gene (**Fig 2A**), with only 12 bp before the next predicted open reading frame (ORF) in the case of SphUPP:SBL2-17, offering an extremely short target. For the other two strains, the gene following *glmS* was co-directional and within 60 bp of the *glmS* stop codon, such that any insertion in the *glmS* 3’ UTR would likely interfere with its transcription. Only SphATUE:S127H113 represented an ideal genetic architecture, with a convergently terminating ORF 697 bp downstream of *glmS*.

The fitness consequences of the *crtI* mutations are beyond the scope of this study. To determine if our *glmS*-targeting insertions were phenotypically neutral, we focused on *S. melonis* FR1, SphUPP:SBL2-17, and SphATUE:S127H113 because these had insertions immediately downstream of *glmS*, though we note that for none of our strains was the insertion free of potential consequences for the downstream coding sequences (**Fig 2A**). We performed competition assays by mixing each tagged strain with its untagged wildtype version in a 1:1 ratio and growing the mixture on 2 week old *A. thaliana* rosettes, with a harvest after one week (**Methods**). We then determined if the ratio of the strains changed as a result of the competition, which would indicate a fitness difference. In a first experiment, we used qPCR to detect the ratio and detected no fitness defects in *S. melonis* FR1 (*glmS*::Tn), but we were not satisfied with the signal to noise ratio (**Supplementary Fig 3**). In a repeat experiment with additional replicates, we counted colony-forming units (CFUs) from each plant sample plated on two media and calculated the ratio of tagged strains (growing on streptomycin and tetracycline) to total bacteria (growing on streptomycin). This revealed a fitness disadvantage in SphUPP:SBL2-17 (*glms*::Tn) and SphUPP:S127H113 (*glms*::Tn) (**Supplementary Fig 3** and **Supplementary Discussion**).

### Suitability of tags for amplicon-based quantification

DNA barcodes should be reproducibly quantifiable by amplicon sequencing over a wide dynamic range. To test ours, we conducted independent trials with unique transposon-specific primers as well as V4 16S rDNA primers that not only amplified our construct (**Fig 1A**) but also each strains’ native 16S rDNA gene. We first mixed DNA from the 10 *glmS-* or *crtI* tagged strains in a balanced, equimolar ratio, and then created a stepwise linear mix and a stepwise exponential mix by manipulating the volume of each DNA stock relative to what was added to the balanced mix (**Fig 3A**). Despite careful measurement, the balanced mix had slightly uneven abundances of the 10 barcodes, and the underrepresented barcodes were consequently underrepresented in the derivative linear and exponential mixes as well. These patterns were captured by both primer sets, for which relative abundances across all mixes was highly correlated (**Fig 3A-B**). We confirmed that both primer sets also accurately captured the expected linear and exponential adjustments; we multiplied the observed balanced abundances by the linear or exponential factors to calculate expected relative abundances in each altered mix, and saw a high correlation with the observed abundances (R^2^=0.96 and R^2^=0.99 respectively, **Fig 3C-D**).

**Figure 3.**
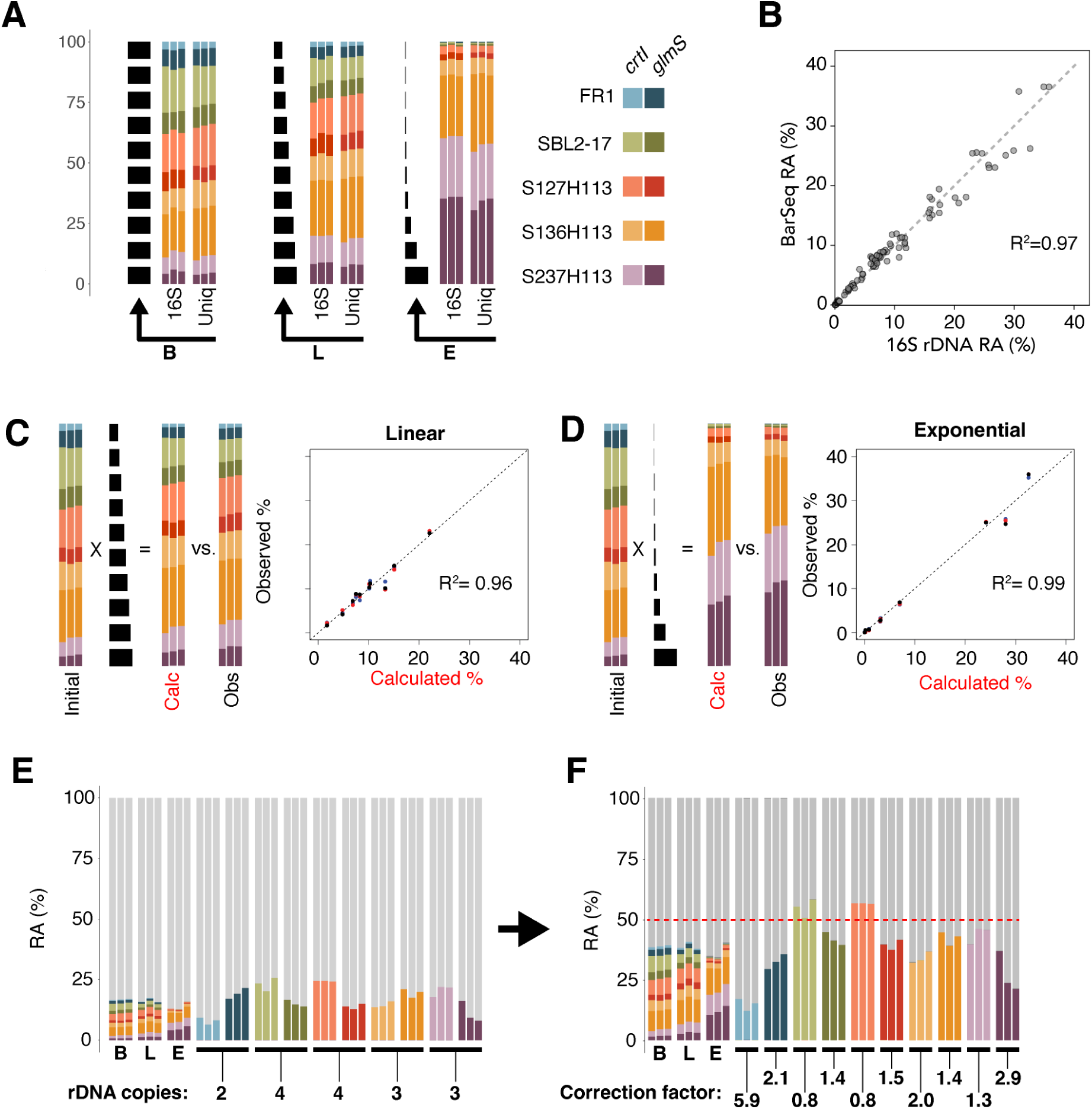
Quantitative validation using DNA mixtures. **A)** Relative barcode abundances (RA) in balanced (B), linearly (L), and exponential (E) mixtures of the 10 tagged strains (colors at right), amplified with either 16S rDNA (16S) or construct-specific primers (Uniq). Each stack of 10 black bars shows strain contributions. **B)** Pearson correlation between barcode abundances measured with 16S rDNA vs. unique primer across all mixes. **C)** Comparison of relative abundances in the linear mix to predicted values obtained after applying the linear multiplication factors to the balanced mix (Cartoon left, correlation right). **D)** Same as (C) for the exponential mix. **E)** Left: Relative abundances of total 16S rDNA amplicons (grey) and barcode amplicons generated with 16S rDNA primers for the three mixes and individually-amplified strains. Genome-derived 16S rDNA copy number shown below. **F)** Same as (E) after correcting for 16S rDNA copy number. The red dotted line (50%) indicates the expected barcode abundance assuming equal amplification efficiency to that of 16S rDNA. The correction factors shown rescale barcode abundances to this expectation.

Our inclusion of V4 16S rDNA priming sites on the tagging construct enables a single 16S rDNA primer pair to detect tagged bacteria and uncharacterized bacteria in a complex sample. However, to be able to compare DNA barcode abundances to 16S rDNA abundances, or to each other, it is important to know how efficiently each DNA barcode is amplified, and how much 16S rDNA signal each tagged strain contributes. We therefore used 16S rDNA primers to sequence amplicons made individually from genomic DNA of each barcoded isolate (**Fig 3E**).

Importantly, the sequence to which the 16S rDNA primers anneal is identical in the native rDNA in all five of our strains, and in our tagging construct as well, so primer annealing should not be a source of bias. After correcting for 16S rDNA copy number in the resulting sequence counts by dividing the 16S rDNA signal by the known number of 16S rDNA copies derived from each genome sequence, it became clear that some variability in barcode amplification existed between barcodes, including between different barcodes within the same strain (**Fig 3F**). Notably, barcodes for *S.melonis* FR1 (*crtI::*Tn) and Sph-ATUE:S237H113 (*glmS*::Tn) showed the lowest amplification efficiency relative to native 16S rDNA (**Fig 3F**), providing a likely explanation also for their underrepresentation in the balanced mix (**Fig 3A**). For each barcode, we then calculated a correction factor by dividing the copy number-normalized 16S rDNA abundance by the mean barcode abundance (**Fig 3F**). Multiplying barcode abundances by the correction factor thus scaled up poorly amplified barcodes and scaled down overly efficient barcodes, such that barcode abundances better reflect strain abundances.

### Quantification of tagged strains in complex environments

For validation of our tagging constructs in a complex environment, we inoculated our *glmS*-targeting tagged strains as a synthetic community on non-sterile *Arabidopsis thaliana* plants. For technical reasons, we decided to also include SphATUE:S136H113 (Tn) and SphATUE:S237H113 (Tn), for which the *glmS*-targeting insertions were considerably off-target. In addition to inoculating the five strains (Mix5), we also created an inoculum of Mix5 combined with a complex uncharacterized community cultured from river water that was enriched for *Sphingomonas* (WaterCom, **Methods**, **Fig 4A**). Interestingly, SphATUE:S136H113 (Tn) was dominant in the Mix5 inoculum, despite the strains having been carefully mixed at equal OD_600_, suggesting this strain may absorb less light at 600 nm, perhaps due to smaller cell size.

**Figure 4.**
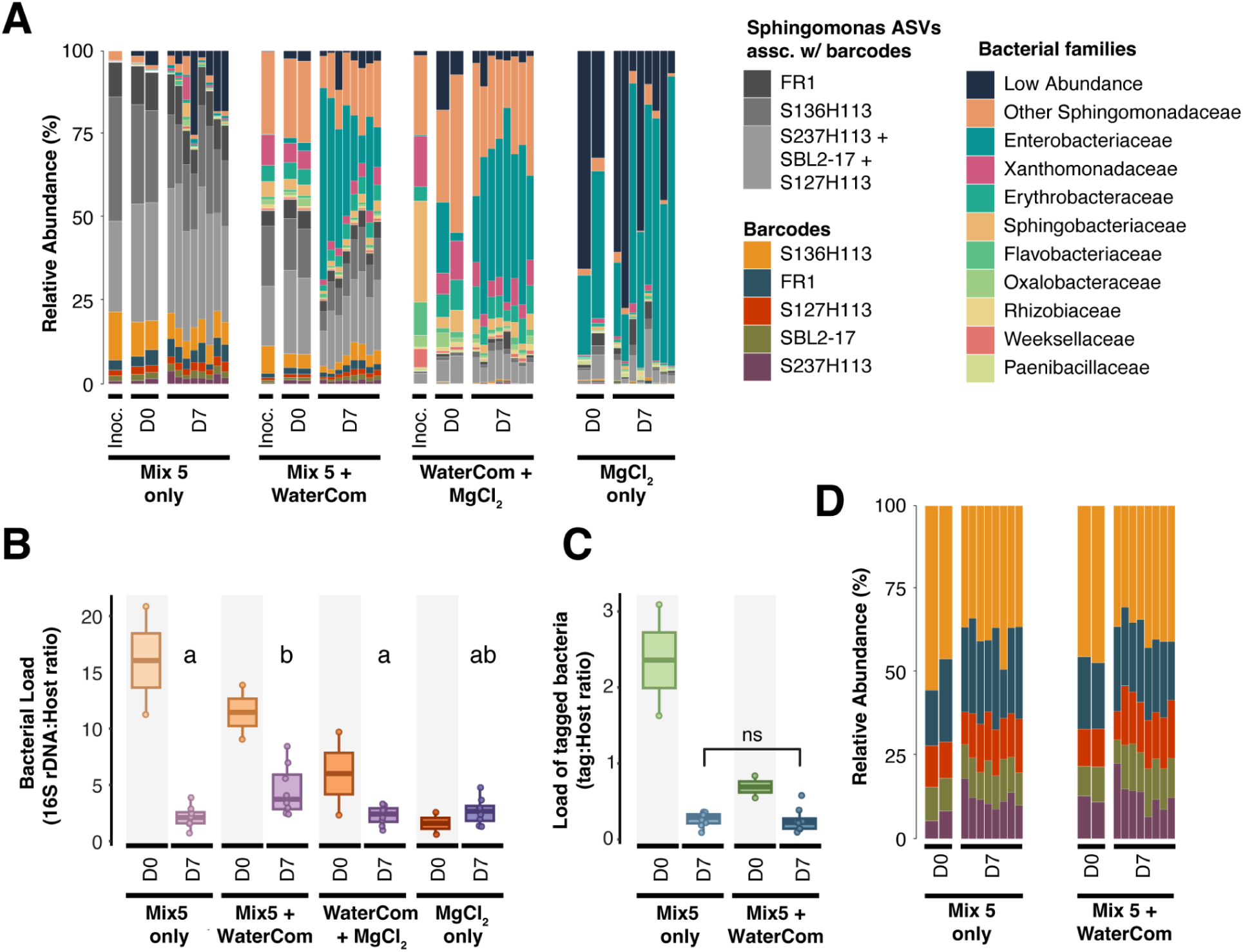
Tagged *Sphingomonas* colonization of plants. **A)** Relative abundances of 16S rDNA amplicons, including both barcodes of the tagging construct and 16S rDNA sequences, for different inoculum conditions. These included an balanced mix of only the barcoded strains (Mix5), a 1:1 mix of Mix5 and an uncharacterized bacterial community isolated from river water (WaterCom), a 1:1 mix of WaterCom with 10 mM MgCl_2_ vehicle, and MgCl_2_ only. The 16S rDNA sequences associated with the barcoded stains are shown in shades of grey (color legend), even if those sequences may have also been already present on the uninoculated plant or in WaterCom. Other bacterial ASVs are classified to the family level (color legend). Inoc, inoculum; D0, day 0 plant sample; D7, day 7 plant sample. **B)** Total bacterial load on plants, determined as the ratio of 16S rDNA (excluding tagging constructs) to the *Gigantea* gene using hamPCR. Statistical differences between groups at D7 were assessed using a Kruskal-Wallis test followed by Dunn’s post hoc test with Benjamini-Hochberg correction for multiple comparisons. Different letters above the boxes indicate significant differences (*p* < 0.05). **C)** Same as (B), but for the load of tagging constructs (excluding all 16S rDNA). Statistical differences between groups at D7 were assessed using a Kruskal-Wallis test. No significant differences were detected (*p* = 0.59; ns). **D)** Relative abundances of the barcodes only for those conditions that included Mix5, following the color legend in (A). In all panels, barcode abundances shown were corrected for technical differences in amplification efficiency (**Methods**).

The high density with which we inoculated plants led to an initial bacterial load that was several fold higher than the ultimate sustainable load that we detected after 7 days (**Fig 4B)**. The load of the barcoded strains alone also stabilized to a similar final level, regardless of the presence of the WaterCom (**Fig 4C)**. While some bacterial families changed in abundance from day 0 to day 7, particularly Enterobacteriaceae which were highly enriched at day 7 in WaterCom-inoculated samples (**Fig 4A**), our barcoded *Sphingomonas* maintained a similar fraction of the community. Moreover, the relative abundance of the barcoded strains among each other was essentially unchanged (**Fig 4D**), suggesting each had similar survival and growth rates in the phyllosphere. Importantly, the 16S rDNA sequences contributed by our tagged strains were present both on uninoculated plants and on plants inoculated only with the WaterCom (**Fig 4A)**, and therefore would have hidden the true abundance of our strains had they not been tagged.

Although three of the five strains in the inoculum shared the same 16S rDNA sequence (**Fig 1D**), *S. melonis* FR1 and SphATUE:S136H113 had unique 16S rDNA, allowing us to again calculate the 16S-to-tag ratio for those strains in the Mix5 inoculum, as we did for individual strains in **Fig 3F**. The ratio we detected this time, especially for *S. melonis* FR1, was considerably different (**Supplementary Fig 4**), likely a result of replication-associated gene dosage [22]. Unlike the stationary phase cultures used in **Fig 3**, the Mix5 inoculum was prepared from bacteria in mid log phase, which may have led to increased rDNA copies. Since replication-associated gene dosage is minimized in stationary phase cultures, the barcode amplification efficiencies as calculated in **Fig 3F** likely more closely reflect the true situation.

However, to rigorously investigate DNA barcode amplification differences, we developed a hamPCR [23] method to compare the abundance of the immediately proximal tetracycline resistance region of the tagging construct to the abundance of the DNA barcode of the same construct (**Supplementary Fig 5**). This “Tet-to-tag” ratio in genomic DNA is always 1:1. Briefly, hamPCR uses template-specific primers for only two PCR cycles to add universal overhangs to two sequence-unrelated templates, removes those primers, and completes the remainder of the exponential PCR with a single pair of primers that anneals to the universal overhangs. This removes primer bias during amplification and preserves the original ratio between the templates, unless those templates amplify differently due to sequence-intrinsic features. These analyses provide support for the barcode correction factors calculated against 16S rDNA in **Fig 3F** (**Supplementary Fig 5 and 6**), and we therefore used these values to scale barcode abundances in **Fig 4**. We note, however, that calculating barcode-specific correction factors from sequence counts of such hamPCR amplicons, rather than from 16S rDNA-primed amplicons, would be preferable moving forward, as it would not be influenced by replication-associated gene dosage.

### An improved universal tagging system

We were unsatisfied with the *glmS* 3’ UTR as a clean integration site in our five strains, due either to small landing pads intolerant of slight targeting errors, or to co-directional downstream genes for which transcription might be interrupted. To see if these potentially problematic features were more widespread in *Sphingomonas*, we investigated genomic structure around *glmS* in 138 non-redundant genomes in our culture collection, ignoring highly-similar genomes differing by less than 1% MASH distance [24]. We recorded the orientation of the next annotated feature and its distance from the stop codon, and whether the 3’ UTR included any other non-coding feature (eg. noncoding RNA). Indeed, in a majority (70%) of cases, the next gene or feature was either < 34 bp from the *glmS* stop codon or was co-directional, and thus would be affected by our transposon (**Fig 5A**). We created an additional sequence resource from which to screen variation by culturing 250 diverse *Sphingomonas* from 25 geographically-separated Arabidopsis rosettes around Uppsala,

**Figure 5.**
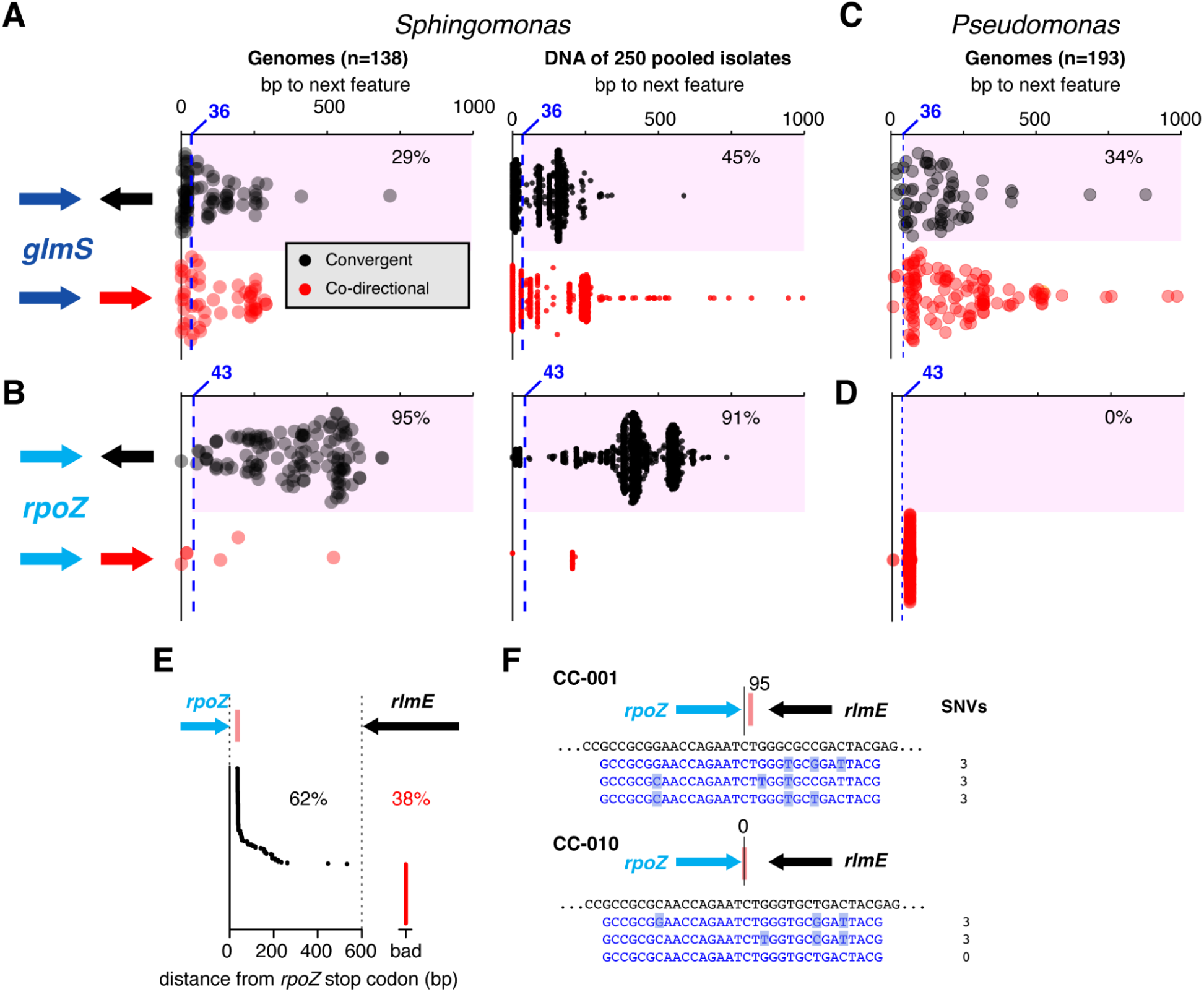
Safer integration loci for high-throughput tagging. **A)** Elements downstream of *glmS* in 138 non-redundant *Sphingomonas* genomes (left) or in a sequenced pool of 250 *Sphingomonas* isolates (right). Zero on the x-axis represents the *glmS* stop codon, and distance along the x-axis represents base pairs until another feature that was convergently terminating (black, above), or co-directional (red, below). The dotted blue vertical line represents the expected transposon insertion location based on the chosen guide family, with its distance from the stop codon indicated. The pink shaded box surrounds acceptable situations for which a convergently terminating feature is downstream of the expected insertion, with the percent of acceptable insertions indicated. **B)** As in (A), but for the region downstream of the *rpoZ* gene. **C-D)** As in (A) and (B), but in a collection of 194 *Pseudomonas* genomes. **E)** tagIMseq mapping for a pool of 200 primary transformants with a guide targeting *rpoZ* in *S. melonis* FR1. The y-axis represents the *rpoZ* stop codon, and the distance to each point along the x-axis represents the distance to an observed transposon insertion. Black dots, and the black percentage, represent acceptable cases in which the insertion precedes the convergently terminating *rlmE* gene, while red dots and the red percentage represent insertions outside of this acceptable range. **F)** Single nucleotide variants between the genomes of isolates CC-001 and CC-010 (black) and the tandem triple *rpoZ* guides (blue) used to transform the pool of 250 *Sphingomonas*.

Sweden (10 isolates per rosette) and sequencing the pool as an unindexed metagenome with long reads. We also saved pool aliquots as a glycerol stock for later bulk culturing (**Methods**). In this sequence resource as well, we annotated *glmS* and downstream features. As with individual genomes, in a majority of cases, insertion downstream of *glmS* appeared problematic (**Fig 5A**).

To search for a more robust insertion location, we used the targetFinder algorithm [25] as well as a pangenome constructed with PPanGGOLiN to search 48 closed *Sphingomonas* genomes for convergently terminating ORFs, requiring a minimum gap size of 50 bp. We found that *rpoZ* (DNA-directed RNA polymerase subunit omega) emerged as a gene flanking such a gap in a majority of strains. A full analysis of the gap revealed that in more than 90% of cases, both in non-redundant *Sphingomonas* genomes and the pooled mix of 250 diverse *Sphingomonas*, the *rpoZ* 3’ UTR provided a likely safe landing pad (**Fig 5B**). We then searched for a guide family in the final 75 bp of *rpoZ*, following the logic described earlier for *glmS*, finding several acceptable candidates (**Supplementary Fig 7**). We extended this analysis to the gamma-Proteobacteria *Pseudomonas* (**Supplementary Table 4**), another common member of the phyllosphere that has been extensively transformed with Tn7 [10,26–28]. Surprisingly, the region downstream of *glmS* was also problematic in many genomes (**Fig 5C**). Further, *rpoZ* did not pose a solution in *Pseudomonas*, with the next element co-directional in all observed cases (**Fig 5D**). However, a new search within *Pseudomonas* revealed that gaps downstream of *pyrD* (quinone-dependent dihydroorotate dehydrogenase), and *fur* (ferric uptake regulator), were excellent genus-wide candidates with convergently terminating gaps in nearly all surveyed strains. Further, as with *Sphingomonas*, there was sufficient gene conservation to find attractive guide families for each gene, with predicted insertions occurring in the safe landing pad (**Supplementary Fig 8**).

To test the *rpoZ* 3’UTR, we synthesized a guide for *S. melonis* FR1 (**Supplementary Table 2**), and generated transformants using pSPIN-LL. To economically map the insertion locations and their associated barcodes, we developed the method <u>Tag</u>mentation transposon Insertion site <u>M</u>apping by <u>Seq</u>uencing (tagIMseq), building on a method by [29], and validated it on a template of pooled DNA from the 10 *glmS*- and *crtI*-tagged strains for which we had previously verified insertion sites by whole genome sequencing (**Fig 2**). We further verified that it worked on a suspension of whole bacterial cells from a single colony of one of those strains (**Methods**, **Supplementary Fig 9** and **10**). Equipped with tagIMseq, we pooled 200 *rpoZ*-targeting transformant colonies and mapped all insertions in the pool, which revealed that just over half of transformants were sufficiently on-target to fall in the acceptable landing pad (**Fig 5E**). To isolate correctly tagged colonies, we further improved our screening pipeline to first use colony PCR with an *rpoZ*-specific primer and a transposon-specific primer to confirm the presence of a putatively on-target genomic insertion as a band visualized by gel electrophoresis. We then used the same colony suspension for tagIMseq for precise transposon mapping and off-target screening for those PCR-positive colonies. In all six cases tested, tagIMseq revealed PCR-positive colonies had a single on-target insertion in the *rpoZ* 3’ UTR (**Supplementary Fig 7**).

We selected an on-target insertion and conducted a similar fitness test as we had used for *glmS*-targeting mutants, and detected no significant fitness effect (**Supplementary Fig 3 and Supplementary Discussion**). We therefore decided to establish feasibility for high-throughput tagging using the *rpoZ* guide family by using it to target our uncharacterized pool of 250 diverse *Sphingomonas*. To prepare the most broadly-useful guides, we extracted *rpoZ* guide sequences from the pool and clustered them at 93% identity (allowing for 2 mismatches in the 32 bp guide) using USEARCH [30]. We then took the representative sequences from the top three most abundant bins and created a triple tandem guide sequence for ligation into pSPIN-LL (**Supplementary Fig 11**). Next, we transformed the pool of 250 strains directly using bacteria washed from the thawed glycerol stock, and plated the conjugation mix on selective media. From our transformants, we analyzed ten colonies by colony PCR to confirm an insertion downstream of *rpoZ*, found two that had a positive signal, and performed tagIMseq on those colonies to identify bases associated with all integration locations. In each case, this revealed a single insertion. We sequenced the genomes of these strains, which further revealed that in both cases the next coding sequence was a convergently terminating Ribosomal RNA large subunit methyltransferase E (*rlmE*) gene (**Supplementary Fig 12**). The two recovered genomes differed from each other with a MASH distance of 0.22 corresponding to variation in approximately 1 in 5 bases [24], and while one of the genomes had an *rpoZ* sequence exactly matching one of the three guides, the other differed from all other guides by 3 bases (**Fig 5F**).

## Discussion

Here we have designed a minimalist tagging construct and targeted it to custom locations in diverse isolates of the genus *Sphingomonas.* Prior to the development of CASTs as biotechnology tools, this would have required homologous recombination, a process which requires inclusion of bacterial genomic sequences on both sides of the insertion construct [18]. While there are efficient tools using linear DNA and short (35-80 bp) regions of homology in *E.coli*, [31], use of homologous regions of 500 bp or more is typical for non-model organisms [18]. The longer the homologous sequence required, the less likely the same construct can be recycled for use in a genetically-distinct organism. This practical difficulty has driven the interest in transposons like Tn7 [32], and the ease of inserting longer pieces of DNA with Tn7 is such that, when faced with a situation for which an organism is missing an attTn7 site or if the native attTn7 site is not tolerated, researchers have sometimes taken the circuitous approach of inserting an attTn7 site at a custom location via homologous recombination, such that the innate Tn7 proteins can recognize it [33,34].

Like the bacterial tagging system “MoBacTags” described by Ordon et al. [7], our constructs include conserved 16S rDNA priming sites that allow direct comparison of barcode abundances to background 16S rDNA abundances in a complex sample. However, our priming sites correspond to the V4 region instead of the V5-V7 region of MoBacTags, and our amplicon is precisely size-matched to V4 16S rDNA amplicons. As with MoBacTags, we observed reproducible barcode-specific effects on the tag-to-16S read count ratios for different tags, even within the same bacterial strain (**Fig 3**) [7,35]. Here we followed the approach of Ordon et al. [7] to use the 16S-to-tag ratio in DNA preps from individual strains to calculate a “correction factor” for the barcode abundance in each strain, but we also provide evidence that this approach carries risks and is likely only valid for DNA from stationary phase cultures for which replication-associated gene dosage effects on rDNA are minimized [22] (**Supplementary Fig 4**). The hamPCR methodology we developed to measure barcode-specific amplification bias (**Supplementary Fig 5 and 6**), which is immune from gene dosage effects, should prove a more robust method to measure correction factors. After applying appropriate correction factors, the DNA barcodes can be analyzed essentially as 16S rDNA amplicons using the same pipeline, enabling strain level resolution in complex microbial and host backgrounds (**Fig 4**).

An important insight from our genomic analysis was that the Tn7 site may be more problematic as a fitness-neutral site than is generally assumed. While there are scattered reports of fitness penalties due to Tn7 integration [10,34], in most cases the genomic architecture or functions of the downstream region is not considered [7], and in many cases fitness tests of the tagged strains are not even conducted. It is encouraging that the *Sphingomonas* Leaf 257 tagged by Tn7 in Daniel et al. [8] apparently had no fitness defects in vitro or in planta, because a co-directional putative bi-functional diguanylate cyclase-phosphodiesterase only 78 bp downstream was likely disrupted. While genes downstream of *glmS* may be dispensible, this is almost certainly context dependent [36]. The gene downstream of *glmS* in *S. melonis* FR1, for example, is a ROK family fructokinase which is important for fructose metabolism. While this might not slow growth in rich media, it could be more problematic in some oligotrophic leaf environments where fructose is a primary carbon source [37]. In any case, it is preferable to avoid disrupting open reading frames and operons, and our results indicate that the region downstream of *rpoZ* has a superior genetic architecture as a neutral site in *Sphingomonas*. While the region downstream of *rpoZ* is unsuitable in *Pseudomonas* (**Fig 5**), we identified promising highly-conserved gaps between convergently terminating genes in *Pseudomonas* using the same approach (**Supplementary Fig 8**). This underscores that it is both necessary and feasible to consider different conserved safe sites depending on the genetic background.

Unlike in *E. coli*, where off-target insertions with *Vch*CASTs are rare exceptions [9], in *Sphingomonas* we recovered off-target insertions in more than ⅓ of events, ranging from a few bases to thousands of kilobases away (**Fig 5E**). Therefore, precise insertion mapping and detection of off-targets is especially important. We developed an efficient and economical transposon mapping by sequencing pipeline, tagIMseq (**Supplementary Fig 9 and 10**).

Remarkably, the method works without a DNA prep, and when sequenced promptly tagIMseq allows decisions to be made about individual colonies before subculturing becomes necessary, providing substantial time savings. In addition to efficient screening, it is likely that CAST technology will continue to improve. Recently, a genetic screen for factors influencing *Vch*CAST transposition uncovered activator and inhibitor genes that are distributed differently among Gram negative bacteria, and contribute to poor efficiency in some species [38]. The authors modified their *Vch*CAST delivery vector to express bacteriophage λ-Red genes (exo, beta, and gam) and were able to significantly increase efficiency; such a system may further improve efficiency in organisms like *Sphingomonas*.

Our successful use of a guide with 5 mismatches (**Fig 1F**, **Fig 2**) demonstrates that limited sequence variation is tolerated. While this tolerance could complicate the avoidance of off-target insertions in a genome, it is an advantage when targeting a unique locus across diverse genomes. We further exploited this tolerance for mismatches when designing three guides for *rpoZ* to broadly target plant-associated *Sphingomonas*, and indeed recovered a correctly-tagged isolate that had three nucleotide variants distinguishing it from the nearest of the three guide sequences (**Fig 5F**, **Supplementary Fig 11**). A large synthetic community of highly similar but readily-distinguishable strains would be a transformative resource for the study of bacterial natural genetic variation, enabling associating unique alleles with bacterial fitness [39]. Further, streamlining the process of disrupting and tagging orthologous gene families across diverse strains extends mechanistic microbial genetics from the model-strain level to the population level.

## Methods

### Creation of pSPIN-LL barcode library

As a transposon payload, the pSPIN-LL plasmid includes 1) a tetracycline resistance gene flanked by flippase (frt) sites, followed by 2) a transcription termination region and a unique 16 bp barcode flanked by both V4 16S rRNA priming sites and unique priming sites. We also modified the pSPIN backbone outside of the payload by adding a synthetic rpsL gene from pVHD [17] that serves as a negative selection marker in most *Sphingomonas* [18]. The plasmid library was constructed as described below.

#### Construction of pSPIN-LL payload

Our construct design was largely ordered directly as a synthetic construct (Integrated DNA Technologies, Coralville, IA, USA), but also involved some cloning intermediates. We describe only an overview of the intermediate processes, as they did not proceed directly to our final construct and ultimately are not important because the final sequence of the construct is published here and the plasmid itself is available upon request. Briefly, our original synthetic fragment included a flippase site, a gentamicin resistance gene, a green fluorescent protein (GFP), another flippase site, and finally a region flanked by useful PCR sites that included transcription terminators and a 16 bp unique barcode. The barcode diversity was created by amplifying the construct with a custom oligo including random bases. We decided to replace the gentamicin and GFP in the original cassette with tetracycline resistance, and we therefore amplified the tetR promoter and tetR gene from pVHD [17] using primers L0129 and L0130, and amplified other parts of the construct we wished to keep and spliced them together into cloning vector pJET1.2 (Thermo Fisher Scientific, Waltham, MA, USA) using the GeneArt Gibson assembly kit (Invitrogen, Carlsbad, CA, USA). We collected hundreds of colonies, each representing unique barcodes, and re-isolated plasmid from the pooled colonies.

#### Adding construct and *rpsL* to pSPIN

The synthetic *rpsL* gene from pAK405 conferring streptomycin sensitivity [18] was amplified using primers L0066 + L0067, and moved by Gibson cloning into a pSPIN backbone that we had been previously cut with FastDigest KpnI (Thermo Fisher Scientific, Waltham, MA, USA). This construct was then used as a template to amplify the region from the 3’ end of the payload site through the newly-added *rpsL* gene, using the primer pair L0077+L0078. In parallel, we amplified the full construct from pJET1.2 using L0075+L0074. Finally, we cut again the original, unmodified pSPIN with FastDigest XhoI and Fastdigest KpnI and gel purified the vector. We finally cloned the insertion construct (L0075+L0074) and the plasmid *rpsL* fragment (L0077+L0078) into pSPIN using the Gibson method to make pSPIN-LL (**Supplementary Fig 1**), and selected transformants on 10 µg/mL tetracycline plates.

#### Generating additional barcode diversity in pSPIN-LL

The first instance of pSPIN-LL construct had around 400 unique barcodes. To generate a more diverse library, we used pSPIN-LL as a template and re-amplified the majority of the payload using primers L0075+L0022, and we amplified the barcode and *rpsL* gene with L0261+L0078. These fragments were cloned by Gibson into original unmodified pSPIN that had been cut with both XhoI and KpnI and gel purified, thus remaking pSPIN-LL such that each colony represents a unique barcode.

### Checking guide RNAs for off-target risks

Annotation-independent off-target analysis was performed using Bowtie2 (v2.5.4) within the framework established by CAST-guide-RNA-tool (https://github.com/sternberglab/CAST-guide-RNA-tool), with slight modifications. While the same computational environment was used for the analysis, a customized Bowtie2 search strategy was used of end-to-end alignment to verify the full 32 bp gRNA sequence, a 5 bp seed length (-L=5), zero allowable mismatches in the seed region (-N=0), and increased search depth (-D=10). Potential off-targets were identified through direct quantification of mismatch counts in the resulting SAM files and visualization performed in RStudio using the ggplot2 [40] package.

### Plasmid DNA Extraction

Plasmids were extracted from *E.coli* by alkaline lysis [41]. For cloning into pSPIN-LL an additional chloroform purification was performed by adding 400 µL chloroform to the resuspended plasmid DNA, vortexing, and mixing the clarified aqueous phase with 1/9 volume 3M Sodium Acetate (NaAc) and 8/9 volume of isopropanol to form a precipitate collected by centrifugation. This pellet was washed with 80% ethanol, and finally resuspended in 500 µL of elution buffer (EB; 10 mM Tris pH 8.0). Purified plasmid DNA was treated with 1µL / 100 µL of RNase A (10 mg/mL, Thermo Fisher Scientific, Waltham, MA, USA), and further cleaned with solid-phase reversible immobilization (SPRI) beads [23] using a 1:1 ratio of beads to DNA. DNA concentration was quantified using a Qubit fluorometer (Thermo Fisher Scientific, Waltham, MA, USA) following the manufacturer’s instructions.

### Inserting guides into pSPIN-LL

Cloning of guide sequences into pSPIN-LL plasmids was performed as described in [42]. The 32 bp target sequence for the CRISPR integrase plus the 2 bp Protospacer adjacent motif (PAM) site was ordered as two complementary oligos that were phosphorylated with T4 Polynucleotide Kinase (Thermo Fisher Scientific, Waltham, MA, USA) to enable ligation and annealed to make a duplex. Primer names used for oligo duplex are detailed in **Supplementary Table 2**. All primer sequences can be found in **Supplementary Table 1**.

### Confirmation of correct guide insertions in *E. coli*

Colonies of transformed *E. coli* that grew on tetracycline selective media were screened by colony PCR using one primer specific to the guide sequence (the one starting with 5’TTCAC…) and another specific to the transposon construct (L0056). See **Supplementary Table 1-2**

### Transformation of *Sphingomonas*

The *E. coli* diaminopimelic acid (DAP) auxotrophic donor strain WM3064 was made electrocompetent as in [43]. A ligated pSPIN-LL plasmid including a guide sequence and the randomly barcoded construct were introduced by pipetting 5 µL of the resuspended ligation reaction into 50 µL of chilled electrocompetent *E. coli* in 1 mm cuvettes. The mixture was pulsed with standard conditions for *E. coli (*25 µF, 200 ohm, 1800 V) using the Gene Pulser Xell (Bio-Rad Laboratories, Hercules, CA, USA). Following the recovery period, the *E. coli* cells were pelleted by centrifugation, resuspended in 60 µL of LB + 0.3 mM DAP (Thermo Fisher Scientific, Waltham, MA, USA) liquid medium, and plated on LB agar plates containing 10 µg/mL tetracycline. The plates were incubated overnight at 37°C, allowing only the bacteria carrying the modified plasmids to grow.

The plasmids were conjugated from the above transformed *E. coli* cells to *Sphingomonas* through a direct cell-to-cell contact mechanism during an overnight co-culturing step. *E. coli* cells carrying the modified pSPIN-LL vectors were grown overnight at 37°C in LB media with 0.3 mM DAP and 10 µg/mL tetracycline. When they reached an OD600 of 1.0, 3 mL of bacterial culture was pelleted, washed via resuspension in fresh LB liquid supplemented with DAP, and resuspended in 500 µL LB media. Similarly, the *Sphingomonas* strains were grown at 28 °C for 2 days in LB media with 100 µg/mL streptomycin, and 3 mL was washed and resuspended in 500 µL of LB. The *E. coli* and *Sphingomonas* were combined, and this 1 mL was again pelleted, resuspended in 60 µL LB liquid with DAP, pipetted as dot on LB agar with DAP but no antibiotic, and incubated at 28°C for 16 hours.

The conjugation mix was then scraped off with a sterile loop, plated onto LB Agar plates with 10 µg/mL tetracycline and 100 µg/mL streptomycin and without DAP to remove *E. coli* cells and select for transformed *Sphingomonas* without pSPIN-LL.

### Confirmation of transformed *Sphingomonas*

#### By colony PCR

Initially, colony PCR was performed only to rule out the continued presence of the pSPIN-LL plasmid using primers L0066 + L0067 specific to the synthetic *rpsL* construct, and to verify the presence of the transposon insertion using unique primers L0116 and L0262. Later, only a primer binding to the guide location (eg. L0427) and a primer binding to the 5’ end of the insertion construct (L0120) was used to reveal colonies with insertions within close range of their insertion.

#### By whole genome sequencing

For each of our five focal *Sphingomonas* strains, DNA from one strain each for the *crtI* and *glmS*-targeting transformations was pooled, and bacterial genome sequencing on the pool of 2 was performed by Plasmidsaurus (Plasmidsaurus Inc., South San Francisco, CA, USA) using Oxford Nanopore Technology with custom analysis and annotation. Reads containing the barcoded transposon construct were identified in the raw fastq file using the following regular expression and its reverse complement. Genomic regions just upstream and downstream of the transposon border were used to confirm that each barcode was associated with a single insertion, and to map that location.

"GCCTTTTTGCGT[ACGT]{16}TTACCGCGGCTGCTG"

### By <u>Tag</u>mentation transposon Insertion site Mapping sequencing (tagIMseq)

Briefly, our method uses Tn5-based tagmentation followed by PCR using an insertion-specific primer and an adapter-specific primer, and is adapted with some additional modifications from [29].

### Template preparation

For re-analyzing the ten *crtI* and *glmS*-targeting mutants as a pool, individual DNAs from each strain were combined. For analyzing *rpoZ*-targeting transformants as pools, colonies from the initial transformation were subcultured once by plating in a grid on a fresh streptomycin and tetracycline plate. After 24-48 hours of growth, all bacteria from the grid were pooled and the pellet was used for DNA extraction as described below. For analyzing individual bacteria, high quality DNA was either prepared from individual isolates, or, for colony tagIMseq, a single colony was resuspended in 14 µL of water and 7 µL was used as template.

### Preparation of assembled Tn5

Tn5 was procured from Protein Production Sweden. Duplexes were prepared by annealing the appropriate oligonucleotides (listed below) in annealing buffer containing 100mM Tris (pH 8.0) and 500 mM NaCl. Duplex A was generated by mixing 9 µL of 100µM Tn5ME_A (L0361) with 9µL of 100µM aminoME_rev (L0376) and 2 µL annealing buffer. Duplex B was prepared similarly using Tn5ME_longB (L0416) and aminoME_rev (L0376). Oligonucleotides were annealed by incubation at 98°C for 3 minutes, followed by cooling to 23°C at a rate of 0.1°C /sec. Duplex A and B were then mixed. Tn5 stock was diluted by mixing 1 µL of stock with 16 µL of dilution buffer containing 50mM Tris-HCl (pH 7.5), 100 mM NaCl, 0.1 mM EDTA, 1 mM DTT, 0.1% Triton X-100, and 50% glycerol. To the diluted Tn5, 3 µL of the mixed duplexes were added, and the mixture was incubated at room temperature for 30 minutes.

### Tagmentation and initial PCR amplification

Both the tagmentation and amplification reactions were performed in a Bio-Rad T100 thermal Cycler (Bio-Rad Laboratories, Hercules, CA, USA). For DNA templates, the tagmentation mixture included 1 µL of DNA (10 ng/µL), 0.2 µL of assembled Tn5, and 2 µL of 5x TAPS-DMF-MgCl [44]₂, and 6.8 µL nuclease-free water for a final volume of 10 µL. For bacterial colony templates, the colony was resuspended in 14 µL water and tagmentation was performed with 7 µL of this resuspension as template, 1 µL assembled Tn5, and 2 µL 5x TAPS-DMF-MgCl_2_. Tagmentation mixtures were incubated at 55°C for 7 minutes, followed by Tn5 inactivation with 2.5 µL of 0.2% SDS and incubation at 55°C for an additional 7 minutes. A first round of PCR was performed using primers annealing to the insertion construct just upstream of the barcode (L0426) and to the transposon-added Nextera overhang (L0417). To denature DNA just prior to PCR amplification and prevent “gap filling” at the Tn5-added adapters, 1.5 µL tagmentation product was mixed with 8.5µL nuclease-free water and denatured at 98°C for 3 minutes. The PCR reagents were then added to a final volume of 25 µL, including 0.5 µM each primer, 5x HF buffer, 200 µM dNTPs, and 0.25 µL Phusion™ Hot Start II DNA Polymerases (Thermo Fisher Scientific, Waltham, MA, USA). The PCR program was set to 98°C for 30 seconds, 30 cycles of 98°C for 30 seconds, 68°C for 15 seconds, and 72°C for 50 seconds, and finally a single extension at 72°C for 5 minutes. PCR products were purified using 0.6x SPRI beads and resuspended in 15 µL nuclease-free water.

### Preparation of library

A second round of PCR was performed using 1 µL of the purified first-round PCR product as template and was amplified with a Nextera P7 index primer (**Supplementary Table 1**) binding to the overhang from primer L0417, and a TruSeq index primer (**Supplementary Table 1**) containing the P5 lllumina adaptor, with each at a final concentration of 0.5 µM. The reaction also contained 5×HF buffer, 200 µM dNTPs, 1.25 µL Phusion DNA Polymerase (Protein Production Sweden, working stock: 60 µg/mL; final concentration: 3 µg/mL), and nuclease-free water to a final volume of 25 µL. The PCR program was set to 98°C for 3 minutes, 18 cycles of 98°C for 30 seconds, 68°C for 15 seconds, and 72°C for 50 seconds, with a final extension at 72°C for 5 minutes. The final PCR products were purified using a double-sided SPRI bead cleanup (0.4x-0.6x), and then DNA concentration was quantified using Qubit.

### Read processing and filtering

R1 reads were concatenated with the reverse-complement of the corresponding R2 reads to generate a single artificial continuous sequence. The combined reads were first filtered to keep only those containing the insertion sequence (GCAGGACGCCCGCCATAAA). (grep -Eo "GCAGGACGCCCGCCATAAA.*"). The second filter came with filtering the remaining reads that carried a 17 bp sequence (GGATGGCCTTTTTGCGT) before the insertion barcode, a 19 bp sequence (TTACCGCGGCTGCTGGCAC) after the barcode, and a 20 bp sequence (ACTTTATGGTTGCATCAACA) at the end of the insertion (grep -Eo "GGATGGCCTTTTTGCGT.{16}TTACCGCGGCTGCTGGCAC.*ACTTTATGGTTGCATCAACA").

In addition, filtered reads were required to contain at least 50 bp of sequence downstream of the insertion end. After filtering, all sequences upstream of and including GGATGGCCTTTTTGCGT, as well as downstream of and including TTACCGCGGCTGCTGGCAC, were excluded to extract the insertion barcode and quantify barcode counts. Reads corresponding to the correct barcodes were then used to extract the 50 bp sequence downstream of the insertion. These potential downstream sequences were arranged from high to low by percentage for each barcode, and the highest one was selected for mapping to the reference genome.

### Bacterial and plant DNA extraction

For bacterial pellets, the final pellets were resuspended in at least 750 μL of DNA lysis buffer containing 10 mM Tris pH 8.0, 10 mM EDTA, 100 mM NaCl, and 1.5% SDS. Especially large pellets (greater than 5 mm from the bottom of the tube) were suspended in proportionally greater volumes of buffer to ensure an efficient lysis, and 750 μL was used for lysis. The suspension was pipetted to a screw cap tube containing a 0.5 mL of a 1:1 mixture of 0.1 mm and 0.5 mm zirconia/silica beads (Bio Spec Products Inc., Bartlesville, USA) and homogenized in a FastPrep 24 (MP Biomedicals, LLC, Solon, OH, USA)instrument at 6 m/s for 1 min, following a subsequent incubation at 55°C for 1 hour. The tubes were centrifuged at 10,000 x g for 5 min, and the supernatant (about 600 μL) was mixed with 200 μL sterile 5 M potassium acetate in a 1.5 mL tube to precipitate the SDS. The tubes were chilled on ice for 10 min, then spun at 14,000 x g for 8 min, and the supernatant was transferred to a new 1.5 mL tube. Then, 360 μL SPRI beads were added to 600 μL of the supernatant (0.6:1 ratio). After mixing and incubating on a magnetic rack, the beads were cleaned with 80% ethanol twice, and DNA was eluted in at least 100 μL EB.

For pure bacteria cultures used to generate closed genomes, an additional RNAse and chloroform cleanup step was added. To the eluted DNA, 1 μL of RNAse A (Thermo Fisher Scientific, Waltham, MA, USA) was added and the mixture was incubated at 37°C for 30 minutes. The DNA was diluted in water to 400 μL, and 400 μL of chloroform was added. The aqueous phase was collected and the DNA was precipitated by mixing it with an equal volume of a solution containing 1 part 3M NaAc and 8 parts isopropanol. The precipitated DNA was pelleted by centrifugation at 14,000 x g for 5 minutes. The pellet was washed twice in 70% ethanol and eluted again in at least 100 μL EB.

For plant material, leaves were added to screw-cap tubes containing 0.5 mL of a 1:1 mixture of 0.1 mm and 0.5 mm zirconia/silica beads (BioSpec Products, Inc., Bartlesville, OK, USA), flash frozen in liquid nitrogen, and homogenized in the FastPrep 24 at 4 m/s for 15 s. Then, 750 μL lysis buffer was added and the extraction proceeded as above.

### Pool of 250 *Sphingomonas* cultivation and sequencing

The 250-member *Sphingomonas* pool was produced by taking 25 *A. thaliana* individuals from around Uppsala, Sweden. Rosettes were rinsed in sterile water to remove adhered soil, and crushed in sterile mortar and pestle. The plant homogenates were resuspended in 0.5X phosphate buffered saline (PBS), and mixed with glycerol (20% v/v) to make culturable freezer stocks. Bacteria were recovered from glycerol by plating on low salt LB media plates with streptomycin (100 μg/mL) and cycloheximide (100 μg/mL). After 72 hours of incubation at room temperature, 10 randomly-chosen colonies from each Arabidopsis individual were picked and passed to a fresh LB plate for even growing, resulting in 250 isolates growing in parallel. After 72 hours of incubation, the 250 isolates were scraped off and homogenized in 10 mM MgCl_2_. The bacterial suspension was adjusted to OD_600_ = 1, mixed with glycerol (20% v/v) and stored at -70°C. An aliquot of the resuspended mix centrifuged and the pellet used for DNA extraction as described above for bacterial pellets. The 250 isolates community DNA was sequenced with long reads on half of a PacBio HiFi Revio flow cell (Pacific Biosciences, Menlo Park, CA, USA) to generate approximately 45 Gb (Novogene, Beijing, China).

### Constructing amplicon libraries

#### Of the DNA barcode using unique primers or 16S rDNA primers

Amplification of the uniquely barcoded region was performed in two rounds of PCR. In the first round, construct-specific primers (L0116 and L0262) were used. PCR reactions were performed in a total volume of 25 µL containing 1× Phusion HF buffer, 0.5 µL of 10 mM dNTP mix, 1.25 µL of each 5 µM primer, 1.25 µL of Phusion DNA polymerase (Protein Production Sweden, working stock: 60 µg/mL; final concentration: 3 µg/mL), 1 µL of template DNA (20 - 60 ng), and nuclease-free water to volume. Between the first and second PCR rounds, amplicons were purified using SPRI beads at a 1:1.1 DNA to SPRI ratio and eluted in 12 µL of nuclease-free water. Subsequently, 10 µL of the purified product was used as a template for the second-round of PCR, which was performed using the same reaction composition with Nextera index primers (**Supplementary Table 1**). The first-round PCR was performed for 15 cycles, and the second-round PCR for 20 cycles, resulting in a total of 35 amplification cycles. Amplification using 16S rDNA primers was performed identically, except that primers L0009 and L0010 were used for the first round of PCR. Following amplification, reactions for each amplicon type were run on a 1% agarose gel to verify a distinct band and for rough quantification of product.

Samples were then pooled based on the relative brightness of each band on the gel, as estimated by eye. The final pool was cleaned with SPRI beads with a 0.9:1 SPRI to DNA ratio to remove primer dimers.

#### by hamPCR

For the inoculation experiments described in **Fig 4**, hamPCR was performed similarly to [23]. Primers L0009 and L0010 were used to tag the 16S rDNA region, and primers L0011 and L0012 were used to tag the *Gigantea* gene. Each reaction contained 5 µL of DNA template (25-250 ng), and the protocol consisted of an initial tagging PCR of 10 cycles using DreamTaq DNA Polymerase (Thermo Fisher Scientific, Waltham, MA, USA), followed by bead cleanup and a second exponential PCR of 25 cycles using Nextera index primers and Phusion DNA polymerase. The final pool of all normalized samples was run on a 2% agarose gel to separate the *Gigantea* and 16S rDNA bands. Both bands were cut out, gel purified with the GeneJET Gel Extraction Kit ((Thermo Fisher Scientific, Waltham, MA, USA), and the GI product was mixed with the 16S rDNA product in a 1:4 ratio to reduce the relative contribution of host amplicon to the pool.

To investigate DNA barcode amplification differences, another hamPCR strategy was used targeting the tetracycline gene and the DNA barcode region within the same barcoding construct (**Supplementary Fig 5 and 6**). A 2-cycle tagging step was performed using primers L0467 and L0472 for the tetracycline gene, and L0471 and L0262 for the barcode region using DreamTaq DNA Polymerase. This was followed by a clean up with 1:1.1 DNA to SPRI beads ratio and a 28-cycle exponential PCR with nextera primers as described above. Amplicons were both visualized on a 2% agarose gel and sequenced, and the ratio of the tetracycline region to the barcode region in each strain was used to determine barcode-specific amplification differences.

### Short read sequencing and amplicon processing

#### Using Illumina MiSeq

For most amplicon samples, different sub-libraries were first diluted to 6 nM, pooled by volume, and then sequenced at 80 pmol with ∼5% PhiX on an Illumina MiSeq i100 series (Illumina, San Diego, CA, USA) using a 25M reagent kit with 300 cycles. Sequencing reads were demultiplexed by the Illumina platform, and only forward reads were used for downstream analysis. Primer sequences were identified and trimmed from their respective datasets using USEARCH v11.0.667 [30], including tagging construct specific primers, primers for the bacterial 16S V4 region, and plant *Gigantea* primers [23]. Reads that did not match the expected primer sequences were discarded. The remaining reads were then quality-filtered with USEARCH to retain only high-quality sequences. All filtered reads were pooled and dereplicated at 100% sequence identity using VSEARCH v2.25.0 [45]. Amplicon sequence variants (ASVs) were inferred as zero-radius operational taxonomic units (ZOTUs) using the UNOISE3 algorithm, which performs both error correction and chimera removal [46]. Taxonomic classification of 16S rDNA ZOTUs was carried out using the SINTAX classifier against the RDP 16S training set v18 (21k sequences) [47,48]. Finally, read counts were mapped to the ZOTUs at a 99% sequence identity threshold using VSEARCH to generate the final abundance tables. For the pure-culture calibration dataset, ZOTU counts of endogenous 16S rDNA were normalized to their known genomic copy numbers (**Fig 3E**). Ratios between copy-number–corrected endogenous 16S rDNA ZOTUs and synthetic tagging construct ZOTUs were used to calculate strain-specific amplification correction factors (**Fig 3F**). Finally, these correction factors were applied to the hamPCR dataset (**Fig 4A**) to account for barcode amplification bias before final relative abundance estimation. Bacterial load was calculated as the ratio of 16S rDNA abundance to *Gigantea* amplicons as in [23]. All downstream normalization, aggregation, and visualization steps were performed in R using the packages reshape [49] and ggplot2 [40].

#### Using Oxford Nanopore

All tagIMseq libraries to verify transposon insertion locations were sequenced by Plasmidsaurus as “Premium PCR Sequencing” using Oxford Nanopore Technology.

### Genomes and annotation

Bacterial genomes used in this study and their source are listed in **Supplementary Table 4**. All bacterial genomes were annotated with BAKTA version 1.11.4 [50] using the full database of AA & DNA sequences and HMM & covariance models hosted at https://zenodo.org/records/14916843. To determine neutral insertion sites in *Pseudomonas*, the NCBI RefSeq [51] annotations were also considered.

### Benign insertion site identification

For *Sphingomonas*, the TargetFinder [25] algorithm was used to identify gaps between convergently transcribed genes in 50 closed *Sphingomonas* genomes from our collection, requiring a minimum gap size of 100 bp. Gens names flanking convergent gaps in multiple genomes were ranked to identify the most highly conserved genes associated with gaps. Candidate gaps were then evaluated in a panel of 138 “non-redundant” *Sphingomonas* genomes. The non-redunant panel was created by using MASH [24] distances to identify genomes from [13] and this work (**Supplementary Table 4**) that were more than 99% similar to each other and randomly removed from each cluster all but a single representative.

For *Pseudomonas*, complete genomes from NCBI RefSeq [51] were similarly compared with MASH and redundant assemblies (more than 99% similar) were removed to create a non-redunant panel of 163. PPanGGOLiN (v2.2.1) [52] was used to create a *Pseudomonas* chromosomal pangenome from these by clustering genes with at least 60% similarity over 80% of the gene into families. Only genes of at least 50 bp occurring in families present in at least 95% of genomes were considered. Only genes bordering co-termination gaps with at least 50 bp before the next gene that were not interrupted by rRNA, tRNA, tmRNA, or ncRNA were considered. To be more conservative, the RefSeq annotations were supplemented with non-coding RNA annotations generated by Bakta (1.8.1).

### Experiments with tagged strains in complex plant environments

To evaluate the tracking efficiency of tagged bacterial strains within complex microbial environments, plant experiments were performed using a synthetic community (Mix5) consisting of five *glmS*-targeting strains: *S. melonis* FR1, SphUPP:SBL2-17, SphATUE:S127H113, SphATUE:S136H113, and SphATUE:S237H113. Each bacterium was grown separately in 50 mL of low-salt LB liquid media in a 200 mL baffled flask until reaching a OD_600_of 0.5 - 1.0, with tetracycline (10 μg/mL) and streptomycin (100 μg/mL). Each culture was then centrifuged at 5000 x g and washed twice in 10 mM MgCl₂ to remove antibiotics, and finally resuspended in 10 mM MgCl₂ and adjusted to OD_600_ = 1. An uncharacterized community of primarily *Sphingomonas* (WaterCom) was isolated from a water sample from the Fyris river (Uppsala, Sweden) on solid LB media with streptomycin (100 μg/mL) and cycloheximide (100 μg/mL) incubated for 7 days. Colonies were washed from the plate with 10 mM MgCl_2_, washed twice via centrifugation and resuspension to remove antibiotics, and adjusted to OD_600_ = 1.

Four inoculum conditions were prepared to assess strain persistence and competitive ability: Mix5 alone; a 1:1 ratio of Mix5 to WaterCom; a 1:1 ratio of WaterCom to 10 mM MgCl₂; and a 10 mM MgCl₂-only negative control . To establish a baseline for community composition, 1 mL of each inoculum mixture was collected before inoculation as “Time-0” samples and pelleted for DNA extraction. Ten individual 3-week old *Arabidopsis thaliana* Col-0 plants were inoculated per condition using a 9cc airbrush paintgun (Biltema, Partille, Sweden) distributing 1mL of inoculum mixture between the ten individuals until the leaf surfaces were uniformly sprayed in two applications allowing them to dry in between. All leaves from two of the inoculated plants were harvested immediately following inoculation as “Time-0 plant” samples. The remaining eight plants per condition were grown for 7 days prior to final harvest. To ensure similar biomass as the Time-0 plant samples, only two mid-sized leaves were collected from each plant at the final harvest. Microbial load on plants was determined using hamPCR [23].

### qPCR to determine fitness penalties due to tagging

To quantify both tagged and untagged bacteria, primers annealing to conserved loci in the single copy rpoZ gene were developed (primers L0443 + L0444), producing a 108 bp amplicon. To quantify only tagged bacteria, primers annealing to the tetracycline resistance gene in our tagging construct were developed (primers L0453 + L0544), producing a 143 bp amplicon. Briefly, each DNA sample containing a mixture of wildtype and tagged bacteria was then amplified in three technical replicates per primer pair, using Taq Univer SYBR Green Supermix (Bio-Rad Laboratories, Hercules, CA, USA) on a Bio-Rad CFX Opus 384 machine. On the same plate, a log10 dilution series of tagged *S. melonis* FR1 *crtI*::miniTn, for which the ratio of tagging construct to *rpoZ* is known to be 1:1, was also amplified. Linear regression was then used on the log10 dilution series for each primer pair to create a standard curve linking Ct value to dilution value. The mean Ct value for the three technical replicates in each sample was interpreted with the correct regression to determine an absolute dilution quantity of each template in each sample. Finally, the absolute quantity of tagging construct was divided by the absolute quantity of *rpoZ*. Each 10 µL qPCR reaction contained 5 µL SYBR mix, 0.25 µL of a working stock (10 µM) of each primer, and 1.5 µL DNA. To reduce the presence of *rpoZ* primer homodimers during reaction setup, the working stock was incubated at 95°C for 2 minutes and cooled on ice to prevent correct annealing, and the reaction was set up on ice and placed into the qPCR machine after it had already reached 95°C.

## Supporting information

Supplementary_Table_1_Oligos

Supplementary_Table_4_Genomes

## Data availability

All sequencing data have been deposited in the European Nucleotide Archive (ENA, https://www.ebi.ac.uk/ena). They can be accessed under project number PRJEB108202. Scripts associated with this work can be found at: https://github.com/LundbergLand

## Author contributions

EMB, HB, TY, MT, and DSL conceived the study. EMB, HB, MT, NN, SH, LCH, and DSL performed inoculation and fitness experiments. EMB, MT, NN, and DSL created mutants. TY developed tagIMseq. EMB, HB, TY, KAG, and DSL analyzed data. EMB and DSL wrote the paper with input from all authors.

## Acknowledgements

We thank Fransiska Liesecke (SLU) for early discussions about guide sequences. We thank Sophia Swartz (UC Berkeley) for discussions about *Vch*CAST machinery and transformation tips. *E. coli* WM3064 was provided by Anne Lanois and Alain Givaudan (Univ. Montpellier). TY was supported in part by the Government Scholarship to Study Abroad (GSSA), Taiwan. Computation was enabled by resources provided by the National Academic Infrastructure for Supercomputing in Sweden (NAISS), partially funded by the Swedish Research Council through grant agreement no. 2022-06725. Tn5 and Phusion enzymes were procured through Protein Production Sweden. This project was supported by NGI OpenLab, Sweden. Funding for this work has been provided by a Wallenberg Academy Fellows grant to DSL (https://kaw.wallenberg.org/en, 2021.0102).

