## Supporting_Information_PDF for "A scalable genomic framework for programmable strain tagging in a diverse bacterial genus"

**Affiliations:**
[1]Department of Plant Biology, Swedish University of Agricultural Sciences, 75007 Uppsala, Sweden

[2]Crop Biotechnics, KU Leuven, 3001 Leuven, Belgium

[3]Universidad Politécnica de Valencia, 46022 Valencia, Spain

*,# = equal contribution

**Correspondence:**
 (D.S.L.)

**This PDF file includes:**

| | |
|---|---|
| Supplementary Figure 1 | 2 |
| Supplementary Table 2 | 3 |
| Supplementary Table 3 | 4 |
| Supplementary Figure 2 | 4 |
| Supplementary Figure 3 | 5 |
| Supplementary Discussion | 6 |
| Supplementary Figure 4 | 7 |
| Supplementary Figure 5 | 8 |
| Supplementary Figure 6 | 9 |
| Supplementary Figure 7 | 10 |
| Supplementary Figure 8 | 11 |
| Supplementary Figure 9 | 12 |
| Supplementary Figure 10 | 13 |
| Supplementary Figure 11 | 14 |
| Supplementary Figure 12 | 15 |

**Other supporting materials for this manuscript included elsewhere are:**

Supplementary Table 1, Supplementary Table 4

### Supplementary Figure 1

| Element | Color | Description |
|---|---|---|
| 1 | | Transposon borders |
| 2 | | flippase (frt) sites |
| 3 | | tetracycline resistance |
| 4 | | unique priming site 1 |
| 5 | | 799R + 806R 16S rRNA gene priming site |
| 6 | | Transcription terminators for tetR (rrnB T1, rrnB T2) |
| 7 | | Random 16-mer barcode |
| 8 | | 515F 16S rRNA gene priming site |
| 9 | | unique priming site 2 |
| 10 | | *rpsL* gene conferring streptomycin sensitivity |

Schematic

```
   1       2       3           2    4     5          6           7      8   9    1      10
--Tn-LB----frt--tet_resistance--frt-- uniq 799R + 806R -terminators- barcodes 515F uniq Tn-RB --rpsL-----
```

Construct full sequence:

```
ACCTCAGGCATTTGTTGTTGATACAACCATAAAATGATAATTACACCCATAAATTGATAATTATCACACCCATAAATTGATATTGCCTCTTCATGGTCTAAACTTCAGTAAGTTT
ACGACATTTTCCTCGAGCGACTCACTATAGGGAGAGCGGCCGCCAGATCTTCCGGATGGCTCGAGTTTTTCAGCAAGATACTGACCCTAGGTGCTCTCTTGAAGTACCTATTCCG
AAGTTCCTATTCTCTAGAAAGTATAGGAACTTCAGAGTTAGTACCTAGATTTAGATGTCTAAAAAGCGGGGTACCGAGGACGCGTCAATTCTCGCGAGACGAAAGGGCCTCGATA
CGCCTATTTTTATAGGTTAATGTCATGATAATAATGGTTTCTTAGCACCCTTTCTCGGTCCTTCAACGTTCCTGACAACGAGCCTCCTTTTCGCCAATCCATCGACAATCACCGC
GAGTCCCTGCTCGAACGCTGCGTCCGGACCGGCTTCGTCGAAGGCGTCTATCGCGGCCCGCAACAGCGGCGAGAGCGGAGCCTGTTCAACGGTGCCGCCGCGCTCGCCGGCATCG
CTGTCGCCGGCCTGCTCCTCAAGCACGGCCCCAACAGTGAAGTAGCTGATTGTCATCAGCGCATTGACGGCGTCCCCGGCCGAAAAACCCGCCTCGCAGAGGAAGCGAAGCTGCG
CGTCGGCCGTTTCCATCTGCGGTGCGCCCGGTCGCGTGCCGGCATGGATGCGCGCGCCATCGCGGTAGGCGAGCAGCGCCTGCCTGAAGCTGCGGGCATTCCCGATCAGAAATGA
GCGCCAGTCGTCGTCGGCTCTCGGCACCGAATGCGTATGATTCTCCGCCAGCATGGCTTCGGCCAGTGCGTCGAGCAGCGCCCGCTTGTTCCTGAAGTGCCAGTAAAGCGCCGGC
TGCTGAACCCCCAACCGTTCCGCCAGTTTGCGTGTCGTCAGACCGTCTACGCCGACCTCGTTCAACAGGTCCAGGGCGGCACGGATCACTGTATTCGGCTGCAACTTTGTCATGA
TTGACACTTTATCACTGATAAACATAATATGTCCACCAACTTATCAGTGATAAAGAATCCGCGCGTTCAATCGGACCAGCGGAGGCTGGTCCGGAGGCCAGACGTGAAACCCAAC
ATACCCCTGATCGTAATTCTGAGCACTGTCGCGCTCGACGCTGTCGGCATCGGCCTGATTATGCCGGTGCTGCCGGGCCTCCTGCGCGATCTGGTTCACTCGAACGACGTCACCG
CCCACTATGGCATTCTGCTGGCGCTGTATGCGTTGGTGCAATTTGCCTGCGCACCTGTGCTGGGCGCGCTGTCGGATCGTTTCGGGCGGCGGCCAATCTTGCTCGTCTCGCTGGC
CGGCGCCACTGTCGACTACGCCATCATGGCGACAGCGCCTTTCCTTTGGGTTCTCTATATCGGGCGGATCGTGGCCGGCATCACCGGGGCGACTGGGGCGGTAGCCGGCGCTTAT
ATTGCCGATATCACTGATGGCGATGAGCGCGCGCGGCACTTCGGCTTCATGAGCGCCTGTTTCGGGTTCGGGATGGTCGCGGGACCTGTGCTCGGTGGGCTGATGGGCGGTTTCT
CCCCCCACGCTCCGTTCTTCGCCGCGGCAGCCTTGAACGGCCTCAATTTCCTGACGGGCTGTTTCCTTTTGCCGGAGTCGCACAAAGGCGAACGCCGGCCGTTACGCCGGGAGGC
TCTCAACCCGCTCGCTTCGTTCCGGTGGGCCCGGGGCATGACCGTCGTCGCCGCCCTGATGGCGGTCTTCTTCATCATGCAACTTGTCGGACAGGTGCCGGCCGCGCTTTGGGTC
ATTTTCGGCGAGGATCGCTTTCACTGGGACGCGACCACGATCGGCATTTCGCTTGCCGCATTTGGCATTCTGCATTCACTCGCCCAGGCAATGATCACCGGCCCTGTAGCCGCCC
GGCTCGGCGAAAGGCGGGCACTCATGCTCGGAATGATTGCCGACGGCACAGGCTACATCCTGCTTGCCTTCGCGACACGGGGATGGATGGCGTTCCCGATCATGGTCCTGCTTGC
TTCGGGTGGCATCGGAATGCCGGCGCTGCAAGCAATGTTGTCCAGGCAGGTGGATGAGGAACGTCAGGGGCAGCTGCAAGGCTCACTGGCGGCGCTCACCAGCCTGACCTCGATC
GTCGGACCCCTCCTCTTCACGGCGATCTATGCGGCTTCTATAACAACGTGGAACGGGTGGCATGGATTGCAGGCGCTGCCCTCTACTTGCTCTGCCTGCCGGCGCTGCGTCGCG
GGCTTTGGAGCGGCGCAGGGCAACGAGCCGATCGCTGATACACATGACATGGACGAGCTGTACAAGTAATAATCTTGAAGTACCTATTCCGAAGTTCCTATTCTCTAGAAAGTAT
AGGAACTTCAGAGGGACTACCAGGGTATCTAATCCTGTTCGTATCGCCTCCCTCGCGCCATCAGGGCAGGCATCAAATAAAACGAAAGGCTCAGTCGAAAGACTGGGCCTTTCG
TTTTATCTGTTGTTTGTCGGTGAACGCTCTCCTGAGTAGGACAAATCCGCCGGGAGCGGATTTGAACGTTGTGAAGCAACGGCCCGGAGGGTGGCGGGCAGGACGCCCGCCATAA
ACTGCCAGGCATCAAACTAAGCGAATTCAGAAGGCCATCCTGACGGATGGCCTTTTTGCGTNNNNNNNNNNNNNNNNTTACCGCGGCTGCTGGCACCTGCAGTAGTTTTGCTGAA
ATACTCGATTCACAAAAATATCAACTTATGGTTGTTTTGTGAGATATCAATATATGGTTGTTTTGTGGTTAAGTTGCTGATTATAAATAATTATTAAATATCACTTTATGGTTGC
ATCAACAGGTACCCCATGGTTGACGGATCGCCGCGAAGCCGCTAACTGCGCCGGCAGATCCATATACGAGGAGGAGGCTTCATGCCGACCATCAACCAGCTGGTCCGGAAGGGGC
GCGACCCGCAGAAGGCGAAGTCGAAGGTCCCGGCGATGGAACAGAACCCCCAGAAGCGCGGCGTGTGCACCCGCGTCTATACCACCACCCCGAAGAAGCCCAACTCGGCGCTGCG
CAAGGTCGCGAAGGTCCGCCTGACCAACAGCCGCGAGGTCATCTCGTACATCCCCGGCGAGGGCCACAACCTACAAGAGCACTCGGTCGTCCTGATCCGCGGCGGCCGGGTCAAG
GACCTGCCGGGCGTCCGCTATCACGTCCTGCGCGGCGTGCTGGACACCCAGGGCGTGAAGGACCGCCGCCAGTCGCGCTCGAAGTATGGCGCGAAGCGCCCGAAGTGATAGGTAC
CGAGCTCGAATTCACTGGCCGTCGTTTTAC
```

**Supplementary Figure 1. Sequence map of tagging construct**
The construct is shown in the "forward" orientation.

### Supplementary Table 2

| Guides *glmS* | PAM (C*) + 32 bp guide sequence | Oligos |
|---|---|---|
| *S. melonis* FR1 | C*GACGTCGACCAGCCGCGCAACCTCGCCAAATC | L0135 + L0136 |
| SphUPP:SBL2-17 | C*GATGTCGACCAGCCCCGCAACCTCGCAAAGTC | L0167 + L0168 |
| Sph-ATUE:S237H113 | C*GACGTCGACCAGCCGCGGAACCTGGCGAAGAG | L0161 + L0162 |
| Sph-ATUE:S127H113 | C*GACGTCGATCAGCCGCGCAACTTGGCGAAGTC | L0165 + L0166 |
| Sph-ATUE:S136H113 | C*GATGTGGATCAGCCGCGCAACCTCGCCAAGTC | L0157 + L0158 |
| SphUPP:SSN-1 | C*GACGTCGACCAGCCGCGCAACCTCGCCAAGTC | L0147 + L0148 |
| Sph-ATUE:S230H113 | C*GACGTCGACCAGCCGCGCAATCTCGCCAAGTC | L0149 + L0150 |
| SphUPP:SATL-3 | C*GATGTCGACCAGCCGCGGAATCTGGCGAAGAG | L0153 + L0154 |

| Guides *crtI* | PAM (C*) + 32 bp guide sequence | Oligos |
|---|---|---|
| *S. melonis* FR1 | C*ATCGTCATCGGATCGGGATTCGGCGGGCTGGC | L0138 + L0139 |
| SphUPP:SBL2-17 | C*ATCGTCATCGGTGCCGGGTTCGGCGGGCTCGC | L0187 + L0188 |
| Sph-ATUE:S237H113 | C*ATTGTCATCGGTGCAGGGTTTGGCGGGCTGGC | L0201 + L0202 |
| Sph-ATUE:S127H113 | C*ATCGTCATCGGCGCAGGGTTTGGCGGGCTGGC | L0189 + L0190 |
| Sph-ATUE:S136H113 | C*ATCGTCATCGGAGCGGGGTTCGGCGGATTGGC | L0183 + L0184 |
| SphUPP:SSN-1 | C*ATCGTCATCGGTGCCGGGTTCGGCGGGCTCGC | L0187 + L0188 |
| Sph-ATUE:S230H113 | C*ATCGTCATCGGTGCAGGGTTCGGTGGGCTGGC | L0179 + L0180 |
| SphUPP:SATL-3 | C*ATAGTCATCGGCGCAGGGTTTGGCGGGCTGGC | L0177 + L0178 |

| Guides *rpoZ* | PAM (C*) + 32 bp guide sequence | Oligos |
|---|---|---|
| *S. melonis* FR1 | C*GCCGCGGAACCAGAATCTCGGCGCCGATTACG | L0407 + L0408 |
| *rpoZ*_tandem_1 | C*GCCGCGGAACCAGAATCTGGGTGCGGATTACG | L0409 + L0410 |
| *rpoZ*_tandem_2 | C*GCCGCGCAACCAGAATCTTGGTGCCGATTACG | L0411 + L0412 |
| *rpoZ*_tandem_3 | C*GCCGCGCAACCAGAATCTGGGTGCTGACTACG | L0413 + L0414 |

### Supplementary Table 3

| **Strain (*glmS*-targeting)** | **Tag barcode** | **Distance after guide to transposon LB** |
|---|---|---|
| *S. melonis* FR1 (*glmS*::Tn) | GTAGTACTATATGATT | 84 |
| SphUPP:SBL2-17 (*glmS*::Tn) | TCGAGGAATGTCTTCT | 137 |
| Sph-ATUE:S127H113 (*glmS*::Tn) | TAAACTGTAATTTAAG | 85 |
| Sph-ATUE:S136H113 (Tn) | GTAAAGGTGGGATCGG | ~900000 |
| Sph-ATUE:S237H113 (Tn) | TTCTATTTTCGGGTCG | ~5800 |

| **Strain (*crtI*-targeting)** | **Tag barcode** | **Distance after guide to transposon LB** |
|---|---|---|
| *S. melonis* FR1 (*crtI*::Tn) | CGACAAGTACTTAAAT | 1376 |
| SphUPP:SBL2-17 (*crtI*::Tn) | ATCTGTACTGTGTTAA | 47 |
| Sph-ATUE:S127H113 (*crtI*::Tn) | ATTGATTACTTGTTTG | 158 |
| Sph-ATUE:S136H113 (*crtI*::Tn) | CTGGGCGATGATGCGT | 52 |
| Sph-ATUE:S237H113 (*crtI*::Tn) | AATATCTATGTTCATT | 990 |

| **Strain (*rpoZ*-targeting)** | **Tag barcode** | **Distance after guide to transposon LB** |
|---|---|---|
| *S. melonis* FR1 (*rpoZ*::Tn) | GTGATTTTATTTGCTC | 45 |
| CC-001 (*rpoZ*::Tn) | TATACCTCCGGGTTCG | 145 |
| CC-010 (*rpoZ*::Tn) | AGGTGGCAGTCCAGCC | 50 |

### Supplementary Figure 2

*glmS*: SphUPP:SBL2-17 guide (top) vs. actual targeted sequence (below)

```
   Guide used:   C*GATGTCGACCAGCCCCGCAACCTCGCAAAGTC
Actual target:   C*GACGTCGATCAGCCGCGCAACCTGGCGAAGTC
                   ** ***** ***** ******** ** *****
```

*crtI*: SphUPP:SBL2-17 guide (top) vs. actual targeted sequence (below)

```
   Guide used:   C*ATCGTCATCGGTGCCGGGTTCGGCGGGCTCGC
Actual target:   C*ATCGTCATCGGCGCAGGGTTTGGCGGGCTGGC
                   *********** ** ***** ******** **
```

**Supplementary Figure 2. Mismatches in guides used for SphUPP:SBL2-17**
Due to an error in the guides used to target *glmS* and *crtI* in SphUPP:SBL2-17, the actual guides used had single nucleotide variants (highlighted blue).

#### Supplementary Figure 3

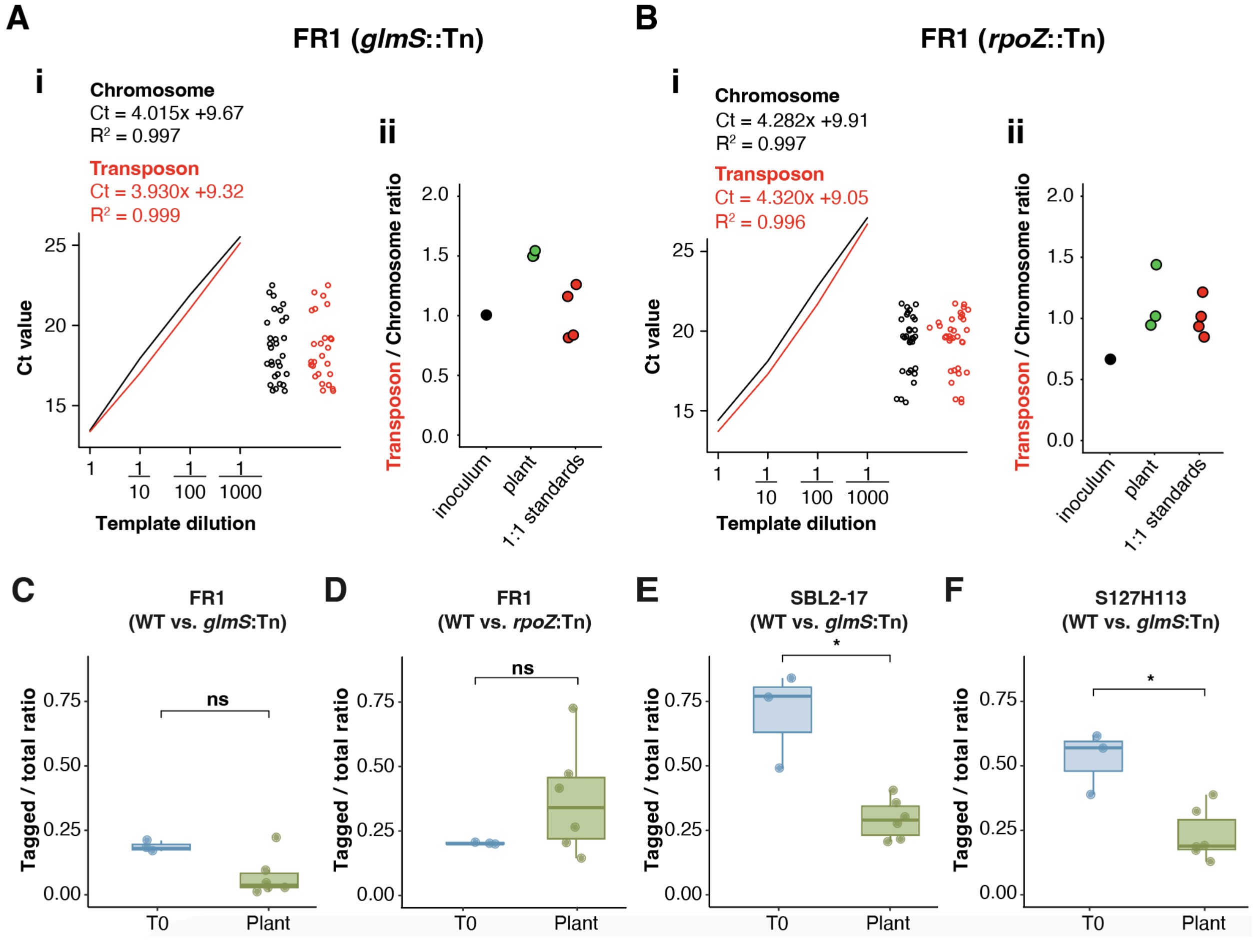


##### Supplementary Figure 3. Fitness of tagged strains compared to wildtype

**A) i:** Standard curves from the chromosome-specific primer pair (black) and transposon-specific primer pair (red) for log10 dilution curves made from a single tagged strain (which has a 1:1 chromosome marker to transposon ratio), with regression equations and $R^2$ shown. Right: actual Ct values from each primer pair for experimental samples in fitness tests comparing FR1 and FR1 (*glmS*::Tn), showing that all experimental Ct values fall within the linear Ct range of the standard curve that was run on the same qPCR plate. **ii:** The calculated ratio of transposon to chromosomal marker in experimental samples. 1:1 standards represent a log 10 dilution series of tagged FR1 DNA used to make the standard curve in (i). **B)** Same as (A), but for FR1 wildtype vs FR1 (*rpoZ*::Tn). **C)** The ratio of tagged colonies (growing on tetracycline + streptomycin) to total colonies (growing on streptomycin) for FR1 tagged downstream of *glmS*, both in the initial mix of both strains before co-culture (T0, blue) and after one week of growth on *A. thaliana* seedlings (Plant, green). The asterisk (*) represents a significant difference ($P < 0.05$ according to a Mann–Whitney U test). **D)** As in C, but for FR1 (*rpoZ*::Tn). **E)** As in C, but for SBL2-17 (*glmS*::Tn) **F)** As in C, but for S127H113 (*glmS*::Tn)

### Supplementary Discussion

For qPCR quantification of fitness tests, the inoculum was prepared as a 1 to 1 mixture of tagged bacteria and wildtype bacteria, and thus had an expected transposon to chromosome ratio of 1 / 2 = 0.5. If wildtype takes over during the experimental competition, the ratio should decrease towards 0, while if the tagged strain is more successful, the ratio should increase towards 1. Ratios above 1 are not possible unless the tagged strain has more than 1 transposon insertion (which we verified was not the case), and thus represent technical artifacts. In these data, pure DNA from tagged bacteria is used in log base 10 dilutions as a standard (filled red points in Supplementary Fig 3A-B), and all are expected to have a transposon to chromosome ratio of 1; deviation from 1 represents noise in the assay.

If one focuses on how the ratio in the competition samples changes relative to ratio in the inoculum prior to co-culture, there is not a clear trend, although if anything, the mutant becomes more highly represented on plants for both FR1 (*glmS*::Tn) and FR1 (*rpoZ*::Tn). While it is possible that two independent transpositions might increase bacterial fitness on plants, the fact that most of the plant ratios are above 1.0, and that even the 1 to 1 initial mixture of FR1 (*glmS*::Tn) had a ratio of 1 ( Supplementary Fig 3A), suggests some kind of technical artifact or noise. There is, at least, no evidence in these data that the tagging construct has made the bacteria *less* fit on plants.

For a higher resolution competitive assay, we conducted a similar fitness experiment. Again we mixed tagged and wildtype bacteria in a 1 to 1 ratio based on $OD_{600}$, and determined the ratio of tagged bacteria as a fraction of all bacteria by plating an equal volume of each sample on two different 90 mm selective plates and counting colonies. We used tetracycline (tet) plus streptomycin (strep) to quantify the tagged bacteria and only strep to quantify total bacteria, and counted a median of more than 70 colonies per plate on the higher selectivity strep+tet plates. These data suggested a significant fitness penalty for all the *glmS*-targeting tagged strains compared to their wildtype version, and only the *rpoZ*-targeting tagged strain appeared unaffected (Supplementary Fig 3C-D), providing additional support for the *rpoZ* 3' UTR as a benign neutral insertion site.

### Supplementary Figure 4

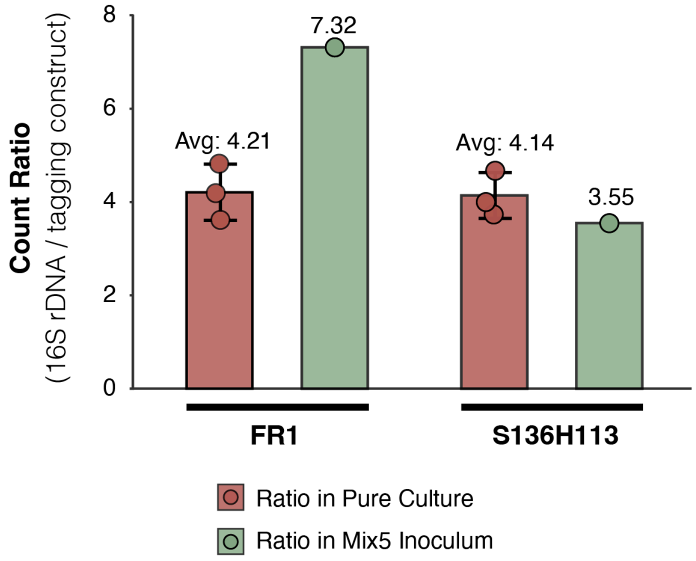


**Supplementary Figure 4. Amplification of tagging construct compared to 16S rDNA in two independent experimental contexts**

Comparison of 16S rDNA to tagging construct read ratios for strains *S. melonis* FR1 (*glmS*::Tn) and SphATUE:S136H113 (*glmS*::Tn) grown in pure culture as in Fig 3 (red, n = 3) versus in a mixed community (Mix5) as shown in Fig 4A (green, n = 1). Analyses were performed with only these strains because they have distinct V4 16S rDNA sequences, making it possible to unambiguously identify both the barcoded tagging construct and the 16S rDNA sequences in the Mix5 community.

### Supplementary Figure 5

hamPCR round 1 tagging primers:

>L0467_tetF
GTCTCGTGGGCTCGGAGATGTGTATAAGAGACAGggctgcaactttgtcatgattg
>L0472_tetR
TCGTCGGCAGCGTCAGATGTGTATAAGAGACAGacgatccgcccgatatagagaacc

>L0471_barcode_region_forward
GTCTCGTGGGCTCGGAGATGTGTATAAGAGACAGcgtatcgcctccctcgcgccat
>L0262_barcode_region_reverse
TCGTCGGCAGCGTCAGATGTGTATAAGAGACAGtgtgaatcgagtatttcagcaaaac

Construct full sequence:

ACCTCAGGCATTTGTTGTTGATACAACCATAAAATGATAATTACACCCATAAATTGATAATTATCACACCCATAAATTGATATTGCCTCTTCATGGTCTAAACTTCAGTAAGTTT
ACGACATTTTCC|TCGAGCGACTCACTATAGGGAGAGCGGCCGCCAGATCTTCCGGATGGCTCGAGTTTTTCAGCAAGATACTGACCCTAGGTGCTCTCTTGAAGTACCTATTCC
GAAGTTCCTATTCTCTAGAAAGTATAGGAACTTCAGAGTTAGTACCTAGATTTAGATGTCTAAAAAGCGGGGTACCGAGGACGCGTCAATTCTCGCGAGACGAAAGGGCCTCGAT
ACGCCTATTTTTATAGGTTAATGTCATGATAATAATGGTTTCTTAGCACCCTTTCTCGGTCCTTCAACGTTCCTGACAACGAGCCTCCTTTTCGCCAATCCATCGACAATCACCG
CGAGTCCCTGCTCGAACGCTGCGTCCGGACCGGCTTCGTCGAAGGCGTCTATCGCGGCCCGCAACAGCGGCGAGAGCGGAGCCTGTTCAACGGTGCCGCCGCGCTCGCCGGCATC
GCTGTCGCCGGCCTGCTCCTCAAGCACGGCCCCAACAGTGAAGTAGCTGATTGTCATCAGCGCATTGACGGCGTCCCCGGCCGAAAAACCCGCCTCGCAGAGGAAGCGAAGCTGC
GCGTCGGCCGTTTCCATCTGCGGTGCGCCCGGTCGCGTGCCGGCATGGATGCGCGCGCCATCGCGGTAGGCGAGCAGCGCCTGCCTGAAGCTGCGGGCATTCCCGATCAGAAATG
AGCGCCAGTCGTCGTCGGCTCTCGGCACCGAATGCGTATGATTCTCCGCCAGCATGGCTTCGGCCAGTGCGTCGAGCAGCGCCCGCTTGTTCCTGAAGTGCCAGTAAAGCGCCGG
CTGCTGAACCCCCAACCGTTCCGCCAGTTTGCGTGTCGTCAGACCGTCTACGCCGACCTCGTTCAACAGGTCCAGGGCGGCACGGATCACTGTATTC**GGCTGCAACTTTGTCAT**
**GATTGACACTTTATCACTGATAAACATAATATGTCCACCAACTTATCAGTGATAAAGAATCCGCGCGTTCAATCGGACCAGCGGAGGCTGGTCCGGAGGCCAGACGTGAAA**
**CCCAACATACCCCTGATCGTAATTCTGAGCACTGTCGCGCTCGACGCTGTCGGCATCGGCCTGATTATGCCGGTGCTGCCGGGCCTCCTGCGCGATCTGGTTCACTCGAAC**
**GACGTCACCGCCCACTATGGCATTCTGCTGGCGCTGTATGCGTTGGTGCAATTTGCCTGCGCACCTGTGCTGGGCGCGCTGTCGGATCGTTTCGGGCGGCGGCCAATCTTG**
**CTCGTCTCGCTGGCCGGCGCCACTGTCGACTACGCCATCATGGCGACAGCGCCTTTCCTTTGGGTTCTCTATATCGGGCGGATCGT**GGCCGGCATCACCGGGGCGACTGGGG
CGGTAGCCGGCGCTTATATTGCCGATATCACTGATGGCGATGAGCGCGCGGCACTTCGGCTTCATGAGCGCCTGTTTCGGGTTCGGGATGGTCGCGGGACCTGTGCTCGGTGG
GCTGATGGGCGGTTTCTCCCCCCACGCTCCGTTCTTCGCCGCGGCAGCCTTGAACGGCCTCAATTTCCTGACGGGCTGTTTCCTTTTGCCGGAGTCGCACAAAGGCGAACGCCGG
CCGTTACGCCGGGAGGCTCTCAACCCGCTCGCTTCGTTCCGGTGGGCCCGGGGCATGACCGTCGTCGCCGCCCTGATGGCGGTCTTCTTCATCATGCAACTTGTCGGACAGGTGC
CGGCCGCGCTTTGGGTCATTTTCGGCGAGGATCGCTTTCACTGGGACGCGACCACGATCGGCATTTCGCTTGCCGCATTTGGCATTCTGCATTCACTCGCCCAGGCAATGATCAC
CGGCCCTGTAGCCGCCCGGCTCGGCGAAAGGCGGGCACTCATGCTCGGAATGATTGCCGACGGCACAGGCTACATCCTGCTTGCCTTCGCGACACGGGGATGGATGGCGTTCCCG
ATCATGGTCCTGCTTGCTTCGGGTGGCATCGGAATGCCGGCGCTGCAAGCAATGTTGTCCAGGCAGGTGGATGAGGAACGTCAGGGGCAGCTGCAAGGCTCACTGGCGGCGCTCA
CCAGCCTGACCTCGATCGTCGGACCCCTCCTCTTCACGGCGATCTATGCGGCTTCTATAACAACGTGGAACGGGTGGGCATGGATTGCAGGCGCTGCCCTCTACTTGCTCTGCCT
GCCGGCGCTGCGTCGCGGGCTTTGGAGCGGCGCAGGGCAACGAGCCGATCGCTGATACACATGACATGGACGAGCTGTACAAGTAATAATCTTGAAGTACCTATTCCGAAGTTCC
TATTCTCTAGAAAGTATAGGAACTTCAGAGGGACTACCAGGGTATCTAATCCTGTT**CGTATCGCCTCCCTCGCGCCATCAGGGCAGGCATCAAATAAAACGAAAGGCTCAGTC**
**GAAAGACTGGGCCTTTCGTTTTATCTGTTGTTTGTCGGTGAACGCTCTCCTGAGTAGGACAAATCCGCCGGGAGCGGATTTGAACGTTGTGAAGCAACGGCCCGGAGGGTG**
**GCGGGCAGGACGCCCGCCATAAACTGCCAGGCATCAAACTAAGCGAATTCAGAAGGCCATCCTGACGGATGGCCTTTTTGCGTNNNNNNNNNNNNNNNNTTACCGCGGCTG**
**CTGGCACCTGCAGTAGTTTTGCTGAAATACTCGATTCACA**AAAATATCAACTTATGGTTGTTTTGTGAGATATCAATATATGGTTGTTTTGTGGTTAAGTTGCTGATTATAAAT
AATTATTAAATATCACTTTATGGTTGCATCAACAGGTAC|CCCATGGTTGACGGATCGCCGCGAAGCCGCTAACTGCGCCGGCAGATCCATATACGAGGAGGAGGCTTCATGCCG
ACCATCAACCAGCTGGTCCGGAAGGGGCGCGACCCGCAGAAGGCGAAGTCGAAGGTCCCGGCGATGGAACAGAACCCCCAGAAGCGCGGCGTGTGCACCCGCGTCTATACCACCA
CCCCGAAGAAGCCCAACTCGGCGCTGCGCAAGGTCGCGAAGGTCCGCCTGACCAACAGCCGCGAGGTCATCTCGTACATCCCCGGCGAGGGCCACAACCTACAAGAGCACTCGGT
CGTCCTGATCCGCGGCGGCCGGGTCAAGGACCTGCCGGGCGTCCGCTATCACGTCCTGCGCGGCGTGCTGGACACCCAGGGCGTGAAGGACCGCCGCCAGTCGCGCTCGAAGTAT
GGCGCGAAGCGCCCGAAGTGATAGGTACCGAGCTCGAATTCACTGGCCGTCGTTTTAC

---

**Supplementary Figure 5. Regions amplified in hamPCR to determine DNA barcode amplification biases.**

The construct sequence is exactly as shown in Supplementary Fig 1, with the tagged region of the tetracycline gene (436 bp) highlighted purple and the tagged region of the DNA barcode amplicon (319 bp) highlighted red.

### Supplementary Figure 6

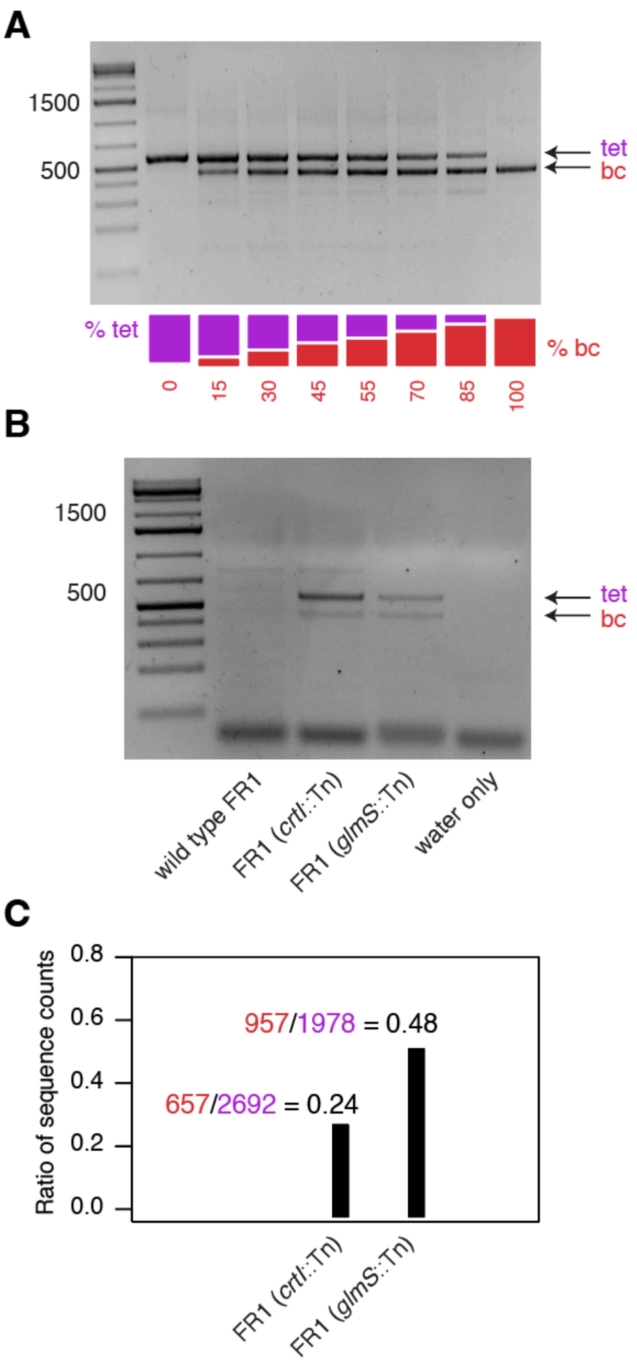


#### Supplementary Figure 6. hamPCR confirms DNA barcode amplification differences

**A)** Tetracycline gene (tet) and DNA barcoded region (bc) hamPCR amplicons visualized on a 2% gel. Amplicons were prepared from mixed templates consisting of different compositions of "tet" and "bc" templates, as shown in the color legend below. The gel band intensities after PCR reflect the initial template abundances, demonstrating that the tagging primers are compatible and assay is quantitative. The sequence of the amplified regions is shown in Supplementary Fig 5. **B)** hamPCR amplicons similar to (A), but prepared from bacterial genomic DNA where the templates are present in a 1:1 ratio, and thus differences in barcode amplification are due to sequence-intrinsic features rather than differences in initial abundance. **C)** Ratios obtained from sequenced amplicons shown in (B) by dividing hamPCR sequence counts mapping to the barcode region (red) by the sequence counts mapping to the tetracycline resistance gene (purple) for *S. melonis* FR1 (*crtI*::Tn) on the left and *S. melonis* FR1 (glmS::Tn) on the right. The amplification of FR1 (*crtI*::Tn) was about half as efficient as FR1 (glmS::Tn downstream), corresponding with the result in Fig 3F.

#### Supplementary Figure 7

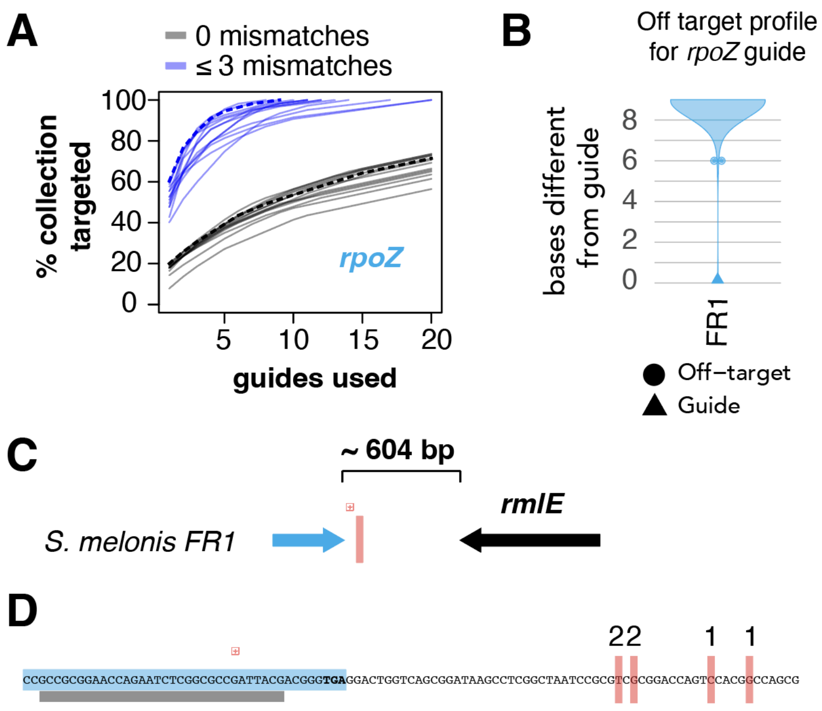


**Supplementary Figure 7. Inserting construct at *rpoZ* in *S. melonis* FR1**

**A)** Number of strains from our culture collection (y-axis) that can in principle be cumulatively targeted by a family of up to 20 guide sequences (x-axis). Each grey or blue line represents a family of 20 sequence-related guides derived from a single conserved PAM site, with grey lines representing the case of perfect alignment to the culture collection, and blue lines allowing for alignments with up to 2 mismatches. The thicker line represents the guide sequence family chosen for subsequent experiments. **B)** Sequence similarity of off-target regions in *S. melonis* FR1 to the *rpoZ* guide designed for that genome as determined via BOWTIE2 alignment. Circles are plotted to represent off-targets only for the most similar off-targets. **C)** Diagram of the region downstream of *rpoZ* in *S. melonis* FR1. The red vertical line represents the region of expected insertion. **D)** Results of tagIMseq insertion mapping on six colony PCR-positive colonies. The 3' end of the *rpoZ* gene is highlighted blue, with the stop codon in bold. The grey bar underlines the bases comprising the guide sequence. Bases replaced by the leftmost border of the insertion construct are highlighted by a vertical red line, with the number of colonies transformed at that location indicated with a number above the line.

#### Supplementary Figure 8

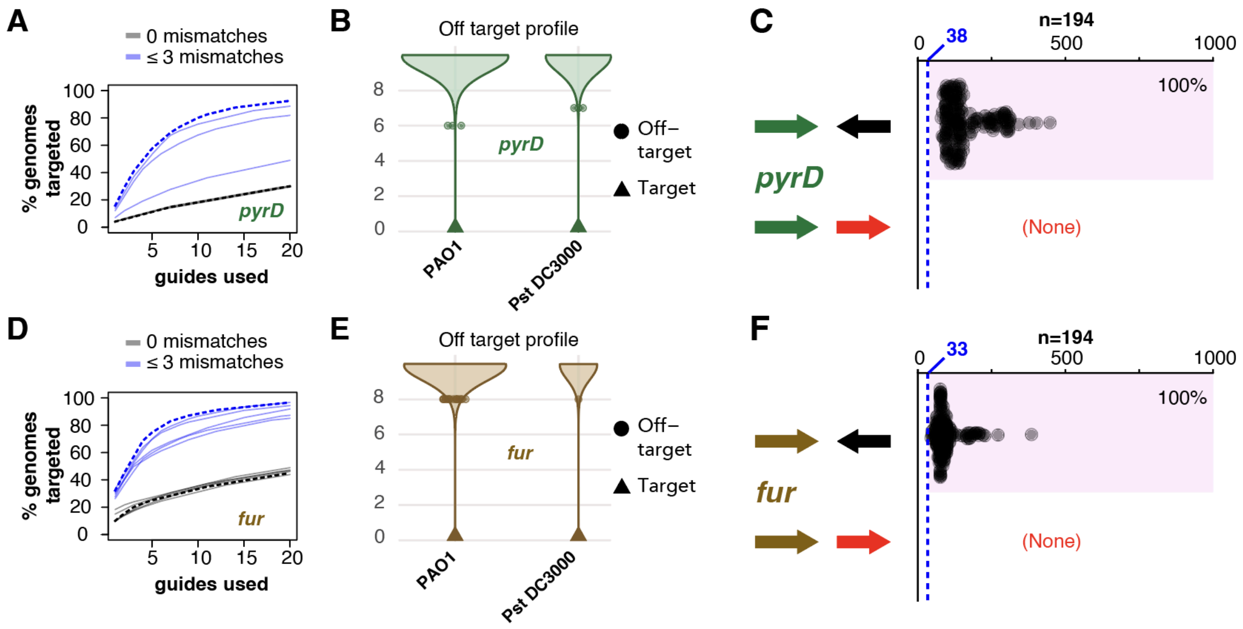


**Supplementary Figure 8. Genus-wide benign insertion sites in *Pseudomonas***
**A)** Percentage of 194 diverse *Pseudomonas* strains (y-axis) that can in principle be cumulatively targeted for transposon insertion downstream of the *pyrD* gene by a family of up to 20 guide sequences (x-axis). Each grey or blue line represents a distinct family of 20 sequence-related guides derived from a single conserved PAM site, with grey lines representing the case of perfect alignment to the culture collection, and blue lines allowing for alignments with up to 3 mismatches. The dotted line represents the hypothesized best guide family. **B)** The most similar off-target regions for a guide in the preferred guide family indicated by the dotted lines in (A) in *P. aeurigenosa* PAO1 (PAO1) and *P. syringae* pv. Tomato DC3000 (Pst DC3000). Circles are plotted to represent discrete off-target events only for the most similar off-targets. **C)** Elements downstream of *pyrD* in 194 diverse *Pseudomonas* genomes. Zero on the x-axis represents the *pyrD* stop codon, with distance along the x-axis representing base pairs until another feature that was convergently terminating (black), or co-directional (red, not represented). The dotted vertical line represents the expected transposon insertion site for the guide family indicated in (A), with its position in base pairs indicated. The pink shaded box surrounds acceptable situations for which a convergently terminating feature is downstream of the expected insertion. The percent of acceptable insertions is indicated as a percentage. **D-F)** As in (A-C), but for the ferric iron uptake transcriptional regulator (*fur*) gene.

### Supplementary Figure 9

## A

##### Duplex A

```
Tn5ME-A, L0361:                     5’-TCGTCGGCAGCGTCAGATGTGTATAAGAGACAG-3’
                                                       |||||||||||||||||||
aminoME_rev, L0376:                         3’-3SpC3-TCTACACATATTCTCTGTC-phosphate 5’
```

##### Duplex B

```
Tn5ME-B_long, L0416:     5’-ACGGCATACGAGATTCGCCTTAGTCTCGTGGGCTCGGAGATGTGTATAAGAGACAG-3’
                                                                  |||||||||||||||||||
aminoME_rev, L0376:                                       3’-3SpC3-TCTACACATATTCTCTGTC-phosphate-5’
```

##### Priming Tn5 adapter in PCR (primes only complement of Duplex B)

```
Round 1 PCR, L0417:                       5’-ACGGCATACGAGATTCGCCTTAG-->
                                             |||||||||||||||||||||||
                                          3’-TGCCGTATGCTCTAAGCGGAATCAGAGCACCCGAGCCTCTACACATATTCTCTGTC...

Round 2 PCR, P7 Nextera Index primer:   5’-CAAGCAGAAGACGGCATACGAGATNNNNNNNNGTCTCGTGGGCTCGG-->
                                                                        |||||||||||||||
                                          3’-TGCCGTATGCTCTAAGCGGAATCAGAGCACCCGAGCCTCTACACATATTCTCTGTC...
```

## B

##### Priming of transposon target in PCR

```
Round 1 PCR, L0425:              5’-CACTCTTTCCCTACACGACGCTCTTCCGATCTGCAGGACGCCCGCCATAAACTG-->
                                                                  ||||||||||||||||||||||
                                                          3’...CGCCCGTCCTGCGGGCGGTATTTGACGGTCC...

Round 2 PCR, P5 Truseq:  5’-AATGATACGGCGACCACCGAGATCTACACNNNNNNNNACACTCTTTCCCTACACGACGC-->
                                                              |||||||||||||||||||||
                                                 3’-GTGAGAAAGGGATGTGCTGCGAGAAGGCTAGACGTCCTGCGGGCGGTATTTGACGGTCC…
```

#### Supplementary Figure 9. tagIMseq adapters and PCR priming

**A)** Alignments of oligos used in Duplex A and Duplex B used to load the Tn5 transposon, and alignments showing how the round 1 and round 2 PCR primers interact with the complement of the long 3’ tail of Duplex B. Green text represents DNA synthesized during the PCR reaction. Red text represents the round 1 primer, and purple text represents the round 2 primer. The N’s represent variable index sequences in the primer. **B)** Alignments showing how the tagging construct is primed during PCR. The N’s represent variable index sequences in the primer.

#### Supplementary Figure 10

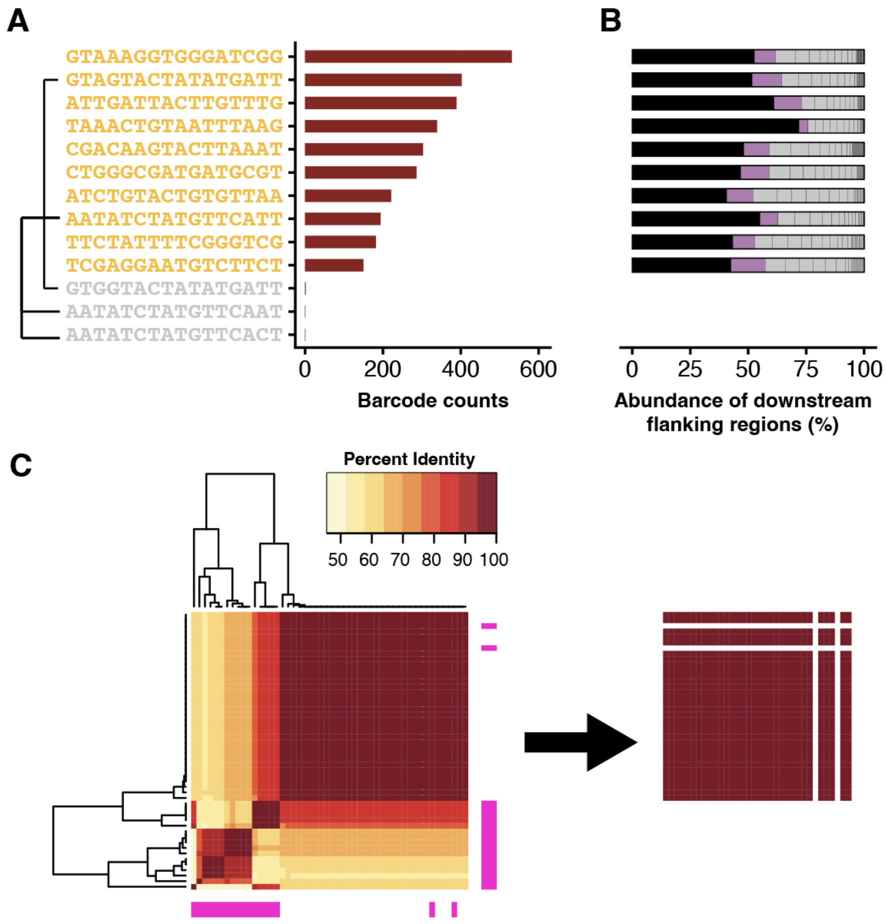


##### Supplementary Figure 10. tagIMseq validation

**A)** Top 13 unique barcode sequences by abundance in the pool of 10 strains, with read counts as indicated. Barcode sequences that are known to be correct from whole genome sequencing are indicated by orange text, and grey text represents other rare barcode sequences that derive from errors in one of the 10 correct barcodes (connecting lines). **B)** The relative abundance of genomic regions downstream of the transposon right border of the tagging construct, as associated with each barcode shown in (A). The known correct genomic region is colored black, the next most abundant associated region colored purple, and all other minor associated regions in grey. In all cases, the purple and grey regions are known to originate from other strains and thus are cases of PCR chimeras. **C)** Relatedness of 50 bp of genomic region downstream of the transposon for colony tagIMseq as sequenced via "Premium PCR" by Plasmidsaurus, with Oxford Nanopore Technology. Sequences matching our transposon construct and containing some downstream sequence are shown at left. Because only a single barcode can integrate into each strain, we required a perfect match to the most abundant barcode sequence. We further eliminated sequences that had our Tn5 adapters in them, and that thus represented less than 50 bp of true genomic sequence. After removing such low quality sequences (pink rows and columns), only highly similar sequences representing a single insertion remained (right).

### Supplementary Figure 11

1) GCCGCGGAACCAGAATCTGGGTGCGGATTACG
2) GCCGCGCAACCAGAATCTTGGTGCCGATTACG
3) GCCGCGCAACCAGAATCTGGGTGCTGACTACG

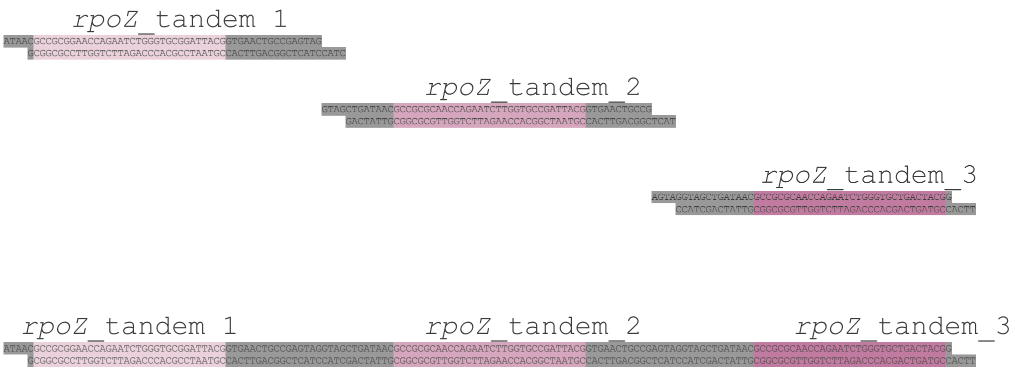


**Supplementary Figure 11. *rpoZ* tandem guide construction**
After extracting the chosen guide location from all sequences in the 250-member *Sphingomonas* metagenome pool and clustering those guides at 93% identity (allowing for two variants between clusters), the three guides shown in blue were the most represented. The guides were constructed into a tandem guide as shown in the diagram.

#### Supplementary Figure 12

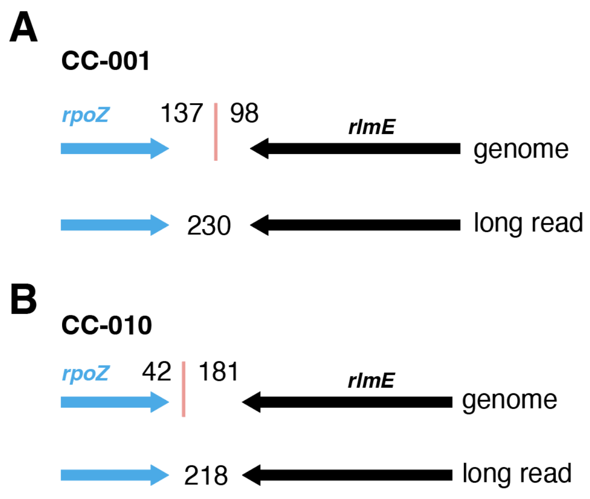


**Supplementary Figure 12. Genomic neighborhood of tagging construct insertions in previously uncharacterized *Sphingomonas*.**
**A)** The whole genome sequence (top) of isolate CC-001 tagged from the pool of 250 Sphingomonas with the triple tandem *rpoZ* guides is juxtaposed with a long read (bottom) obtained from sequencing the pool of 250. The genome sequence and long read have identical sequences except for the transposon insertion in the genome, represented by the vertical red bar. The distance in base pairs from the stop codon of *rpoZ* to the transposon left border and from the transposon right border to the stop codon for *rlmE* are indicated for the genome sequence, and the distance from the *rpoZ* stop codon to the *rlmE* stop codon is shown for the long read. **B)** Same as (A) for isolate CC-010.
